# Improved HIV-1 drug resistance mutation prediction using quasispecies reconstruction supported analysis

**DOI:** 10.1101/2021.05.24.445423

**Authors:** Jyoti Sutar, Shilpa Bhowmick, Varsha Padwal, Vidya Nagar, Priya Patil, Vainav Patel, Atmaram Bandivdekar

## Abstract

Accurate and sensitive approaches to detect HIV-1 drug resistance mutations (DRMs) are indispensable for the paradigm of ‘treatment as prevention’. While HIV-1 proviral DNA allows sensitive high throughput sequencing (HTS)-based DRM detection, its applicability is limited by presence of defective genomes. This study demonstrates application of quasispecies reconstruction algorithms (QRAs) to improve DRM detection sensitivity from proviral DNA. A robust benchmarking of 5 QRAs was performed with 2 distinct experimental control-datasets including a stringent, novel control: DCPM, simulating *in-vivo* variant distribution (0.08%-86.5%). Selected QRA was further evaluated for its ability to differentiate DRMs from hypermutated sequences using an *in-silico* control. PredictHaplo outperformed all others in terms of precision and was selected for further analysis. Near full-genome HTS was performed on proviral DNA from 20 HIV-1C infected individuals, at different stages of ART, from Mumbai, India. DRM detection was performed through residue-wise variation analysis and implementation of QRAs. Both analyses were highly concordant for DRM frequencies >10% (spearman r=0.91, p<0.0001). Phylogenetic association in HTS datasets with shared transmission history could also be demonstrated by PredictHaplo. This study highlights utility of QRAs as an adjunct to traditional residue-wise variation-based DRM detection leading to optimal personalized ART as well as better disease management.

## 1. Introduction

A successful curative strategy for HIV infection has remained elusive due to antigenic variability introduced by the poor proofreading activity of its reverse transcriptase enzyme which affects viral infectivity, response to treatment as well as host immune responses [1]. Antiretroviral therapy (ART) is the only significant treatment modality available for clinical management of HIV infection [2]. To achieve the 90:90:90 treatment goal set by the Joint United Nations Programme on HIV/AIDS, the ART program of National AIDS Control Organization of India, second largest in the world, recently adopted ‘test and treat approach’ [3]. Consequently, with further scale-up of ART [4], at least 2.05 million PLHIV are expected to be receiving treatment by the year 2024 in India [5,6]. However, poor adherence and loss to follow up, conducive for emergence of drug resistance, remain major challenges of the ART program not only in India but also globally [7–9].

High throughput sequencing (HTS) affords unprecedented detection sensitivity in diagnostic pathology [10–12] including HIV DRM prediction [13,14]. While most DRM testing is performed on Viral RNA isolated from blood plasma, recent reports have indicated no difference between DRM predictions based on plasma viremia and peripheral blood mononuclear cells derived proviral DNAs. Rather, proviral DNA is believed to be a richer source of information of viral repertoire as it archives not only present but also DRM harboring quasispecies from the past that may emerge in future causing unexpected accelerated therapy failure [15,16]. Proviral DNA also allows characterization of viral quasispecies from individuals with undetectable plasma viremia [17]. Confounding factors however, include non-replicating viral genomes considered to be phylogenetic artefacts whose inclusion would decrease the sensitivity/accuracy of the analysis and mislead clinical interpretation. Furthermore, residue wise DRM analysis, in contrast to that of extended viral sequences (quasispecies) cannot describe allelic or non-allelic mutation combinations and interactions [18]. Various quasispecies reconstruction algorithms (QRAs), addressing this lacuna have been reported since 2011, but their clinical application has been impaired due to lack of validation standards, complicated implementation and advanced computational requirements [19]. Here, applying sanger sequencing as well as HTS supported by QRAs and a novel control ‘Distributed Concentration Plasmid Mix’ (DCPM), improved prediction of archived DRMs is demonstrated in HIV-1C infected individuals at different stages of ART.

## 2. Materials and Methods

### 2.1. Participant Recruitment and estimation of clinical parameters

HIV-1 clade C infected individuals were recruited with informed consent, from J.J. group of Hospitals, Mumbai following approval of Institutional Ethics Committees. Whole blood samples (10mL) were collected from 32 infected participants (D1-D29) at different stages of ART. Participants were categorized into four groups: ART Naive (**AN**, N=11); First line ART responding (**FLAR**, N=6); First line ART failing (**FLAF**, N=5) and Second line ART receiving i.e. protease inhibitor exposed individuals (**SLAR**, N=7). ART failure was defined per the WHO guidelines as fall of CD4 count to baseline count at the time of initiation of current therapy or plasma viral load >1000 copies/mL after at least 6 months on therapy (Table 1). Detailed description of various clinical parameters such as pre therapy baseline CD4 count, peak CD4 count with therapy, current and past ART combinations received and their duration have been provided in supplementary file 1. CD4 count was estimated by flow cytometry (FACS Calibur, Becton Dickinson, USA). Plasma viral load was estimated using MagNA pure automated nucleic acids isolation system followed by COBAS® Taqman HIV-1 V2.0 assay (Roche, Germany), as per manufacturer’s instructions.

**Table 1.**

| Designation | Age/Sex | Last known<br>CD4 Count<br>(cells/mm<sup>3</sup>) | Last known<br>Viral load<br>(copies/mL) | Category | Sequenced<br>by |
| --- | --- | --- | --- | --- | --- |
| D1 | 40/M | 162 | 304046 | FLAF | SS, HTS |
| D2 | 40/M | 372 | 184723 | FLAF | SS, HTS |
| D3 | 55/M | 124 | 34092 | FLAF | SS, HTS |
| D4 | 48/F | 269 | 7293 | FLAF | SS, HTS |
| D5 | 36/F | 179 | 29194 | FLAF | SS, HTS |
| D6 | 48/M | 414 | <34 | SLAR | SS, HTS |
| D7 | 51/M | 194 | 97 | SLAR | SS, HTS |
| D8 | 45/M | 185 | 848 | SLAR | SS, HTS |
| D9 | 45/F | 202 | 132788 | SLAR | SS, HTS |
| D10 | 42/F | 1684 | TND | SLAR | SS, HTS |
| D11 | 59/M | 194 | TND | SLAR | SS |
| D12 | 32/M | 118 | <34 | SLAR | SS |
| D13 | 37/F | 1061 | TND | FLAR | SS, HTS |
| D14 | 43/M | 714 | <34 | FLAR | SS, HTS |
| D15 | 45/F | 424 | <34 | FLAR | SS, HTS |
| D16 | 45/M | 864 | 457 | FLAR | SS, HTS |
| D17 | 9/M | 1324 | TND | FLAR | SS, HTS |
| D18 | 35/F | 444 | TND | FLAR | SS |
| D19 | 28/M | 534 | 197054 | AN | SS, HTS |
| D20 | 45/M | 612 | 6585 | AN | SS, HTS |
| D21 | 40/F | 440 | 2749 | AN | SS, HTS |
| D22 | 43/M | 527 | 861484 | AN | HTS |
| D23 | 35/M | 230 | 101032 | AN | SS, HTS |
| D24 | 25/F | 510 | 1370 | AN | SS |
| D25 | 36/M | 418 | 32851 | AN | SS |
| D26 | 34/F | 548 | 3456 | AN | SS |
| D27 | 37/F | 536 | TND | AN | SS |
| D28 | 53/F | 762 | 32694 | AN | SS |
| D29 | 34/F | 715 | 668998 | AN | SS |

**Footnote:** FLAF: First line ART failing, SLAR: Second line ART receiving, FLAR: First line ART receiving, AN: ART naive, TND: Target not detected, NA: Not applicable, SS: Sanger Sequencing, HTS: High throughput sequencing

### 2.3. DNA isolation and PCR amplification

Genomic DNA was extracted from peripheral blood mononuclear cells (PBMC) (∼5 × 10^6^) by QIAamp blood DNA mini kit (Qiagen, USA) per manufacturer’s instruction.

#### Protease (PR), Reverse Transcriptase (RT) and Integrase (IN) amplification

PR (HXB2: 2147-2639), RT (HXB2: 2511-3351) and IN (HXB2: 4343-5092) regions were amplified with a nested PCR approach. This strategy was followed for datasets indicated with SS (N=28) in Table 1.

#### Near full length genome (NFLG) amplification

NFLG of HIV-1 was amplified with 3 overlapping fragments covering gag (2570 bp), pol (4089 bp) and env (3133 bp) regions of HIV-1 genome (HXB2: 769-9089) with a nested PCR approach. This strategy was followed for datasets indicated with HTS in Table 1 (N=20).

Details of both amplification strategies along with primer sequences have been provided in Supplementary file 2.

Final PCR products were observed under UV gel documentation system. Equal volumes of PCR products amplified in three independent reactions were pooled to partially mitigate amplification bias and purified with Nucleospin PCR gel purification kit (Machery Nagel) per manufacturer’s instructions.

### 2.4. Control data

#### Experimental error control (EEC)

NFLG amplification was performed on plasmid pIndieC1 [20] (GenBank: AB023804.1) to establish thresholds for errors introduced in amplification and sequencing procedures.

#### Distributed Concentration plasmid mix (DCPM)

DCPM contains five subtype C plasmids with 90-95 percent identity mixed at concentrations of 10ng:∼86.5% (p93IN905)[21]□, 1ng:∼8.65% (p98IN012.14)[22], 500pg:∼4.32% (p94IN476.104)[22], 50pg:∼0.432% (p93IN999) and 10pg:∼0.086% (pIndieC1). Five microliters of this mixture were amplified with NFLG protocol (Supplementary file 3).

#### *In silico* NFLG Control (ISNC)

*In silico* NFLG sequence dataset of 1 million reads was generated with the tool InSilicoseq [23] using HiSeq 2000 profile. Four distinct variants of pIndieC1 sequence containing various combinations of 4 drug resistance associated mutations each for PR, RT and IN were included along with unmutated pIndieC1 as the template at an abundance range from 2% to 50% (supplementary file 4). One of these variant reference sequences was hypermutated and included internal stop codons as well as a unique drug resistance mutation.

### 2.5. Sequencing

idirectional Sanger Sequencing was performed commercially (Eurofins, India) on ABI 3730XL sequencer for PR, RT and IN amplicons. High throughput sequencing was performed commercially (Interpretomics, India) on TrueSeq Nano DNA libraries of NFLG amplicons generated from 2 control and 20 clinical samples on Illumina Hiseq 2000 platform to obtain ‘paired read’ data (read length 101 bases).

### 2.6. Data analysis

#### Sanger Sequencing

Electropherograms obtained post sequencing were inspected and edited with ABI sequence scanner V1 (Applied biosystems) followed by contigs generation with cap3 implementation in BioEdit v7.25[24]. Quality assessment and hypermutations analysis was performed with QC tools available on LANL HIV database [25]. Single nucleotide mutations were detected in electropherograms through NovoSNP [26]

#### High throughput sequencing

Consensus sequences were generated for each sample using VICUNA v1.3 [27]. Quality filtered reads were aligned to their respective consensus sequences using Mosaik v2.2.3 [28]. Variant calling was performed with V-Phaser 2.0 and the output was analyzed with Vprofiler to generate amino acid frequency tables [29,30]. Quasispecies reconstruction (QR) was performed with tools ShoRAH v1.1.3 [31,32], QuRe v0.99971 [33], PredictHaplo v0.4 [34], QsdpR v3.2 [35] and ViQuaS v1.3 [36] for DCPM dataset as well as Five virus mix dataset (SRA: SRR961514) [37] for 17 ORFs of the virus. The data generated was compared with the source viral/plasmid sequences and assessed for recall [(No. of Reconstructed quasispecies closest to the source sequences i.e. accepted quasispecies /expected no of quasispecies) x 100], precision [(No of accepted quasispecies/Total n. of quasispecies generated by the tool) x 100] and processing time: Total duration of QR of PR, RT and IN genes on an intel i7 4790 @ 3.60 GHz (8 threads) processor with 20GB RAM (supplementary file 3). Following this extensive benchmarking, selected QRA was then tested for its ability to detect DRMs using the control dataset INSC. For all the clinical datasets, QR analysis was performed for PR, RT, IN by PredictHaplo. Phylogenetic analyses were further performed for the PR, RT and IN quasispecies with Maximum likelihood (ML) method for sequence data set using PhyML 3.0 [38]□. The best-fit Nucleotide substitution model was predicted by jModelTest2 [39]□. The model selected under Bayesian Information Criterion (BIC) ranking was ‘General time reversible’ (GTR) nucleotide substitution model with a gamma distribution of rates (+G) for all 3 regions. The robustness of the ML tree was further investigated with non-parametric bootstrap analysis available in PhyML with 100 pseudo-replicates. The tree was manually edited for presentation using FigTree v 1.4.0 [40]. Detailed analysis commands are provided in supplementary file 5.

#### Analysis of DRMs

DRM detection was performed with Contig sequences obtained through SS (PR, RT and IN genes) or residue-wise amino acid mutations(*pol*)/quasispecies obtained through HTS using Stanford HIVdb DRM prediction system v8.6.1 with sierrapy utility v2.2.8 [41–43]. DRM datasets thus generated were used to generate heatmaps using the ‘pheatmap’ package of R statistical computing software (v3.4.4) and R studio v1.1.456 [44–46] and Circos v0.69 [47]. Statistical analyses were performed using GraphPad Prism version 5.01 for Windows, (GraphPad Software, USA) and R statistical computing software (v3.4.4) and R studio v1.1.456 [44–46].

### 2.7. Data availability

Raw sequence data is available in Genbank: MH818225-MH818317 and NCBI Bioprojects PRJNA493619 (SRA: SRP162802) and PRJNA493735 (SRA: SRP162818).

## 3. Results

### 3.1. Clinical Features

CD4 counts as well as plasma viremia of the four groups, AN, FLAR, FLAF and SLAR (Table 1) were reflective of therapy status as expected as shown in Figure 1A and B. FLAF individuals had lower CD4 counts compared to AN and FLAR groups and increased viral load indicative of therapy failure. The SLAR group including both therapy responding and failing individuals, overall, exhibited reduced viremia compared to FLAF individuals. However, CD4 counts indicative of rebound, seen in FLAR individuals, were not observed in this group in spite of extended therapy duration (Supplementary file 1). All ART regimens lacked integrase inhibitors and included lamivudine, the most commonly prescribed NRTI under the national ART program of India[48]. Groups FLAF and FLAR received RT inhibitors (RTI) based first line therapy, while the SLAR group received second line therapy consisting of PR inhibitor (PrI) in addition. Therapy failure in recruited participants may be ascribed to self-reported lack of adherence as per their clinical records. Participant D10 reported toxicity (severe rash) to the 1^st^ line regimen as the reason for being switched onto PrI based second line therapy. Of note, the study participants included individuals with shared transmission history. These were a family wherein HIV was transmitted from D16 to D24 heterosexually and further to D17 by mother to child transmission. Also, heterosexual transmission history was recorded from participant D14 to D13.

**Figure 1.**
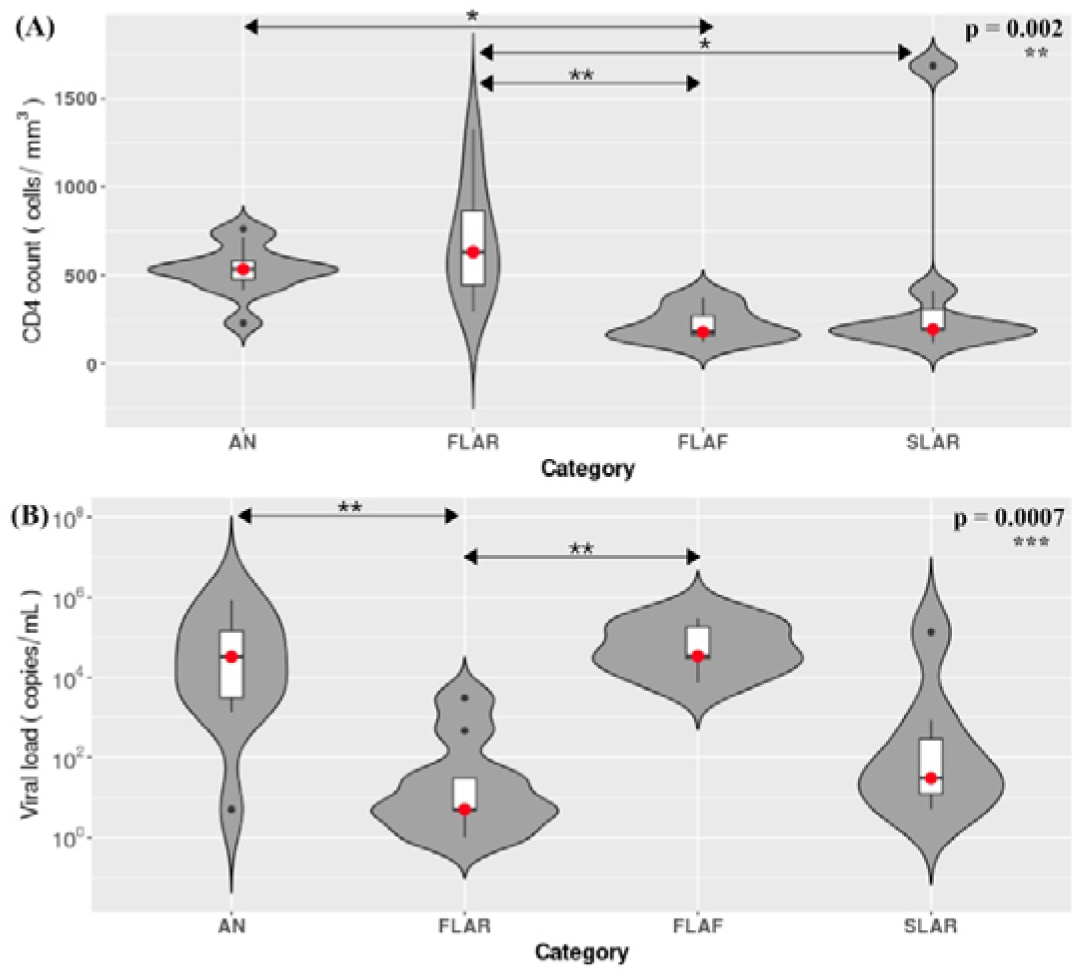
Participant Characteristics: Group wise violin plots for CD4 count (A) and plasma viremia (B). The bar plot in the center of the violin plot denotes 25-75% of the range while the dot on the bar plot indicates the median. Statistical comparisons were made between the groups by One Way ANOVA followed by Dunn’s multiple comparison test and found to be significantly different (A) p=0.002 and (B) p=0.0022

### 3.2. DRM analysis (Residue-wise)

Residue-wise DRM analysis was performed by Sanger sequencing for 28 participants (D1-S29, except D22) and by HTS for 20 participants (D1-D10, D13-D17, D19-D23:Table1) leading to generation of 19 matched data sets. Sanger sequencing-based interpretation described in Figure 2 provided a uni-dimensional, qualitative depiction of drug responses, i.e., presence or absence of DRM most likely present at the highest frequency. HTS analysis, in contrast, allowed quantification of DRMs (Figure 3) present in each sample at frequencies >0.06% based on EEC. The Stanford drug resistance prediction algorithm (HIVDb) does not account for these frequencies in scoring and subsequent interpretation. Therefore, a circos heatmap with frequency weighted color scheme was developed to display this data (Figure 3). Circos heatmap for matched sanger datasets is provided in Supplementary file 6. Furthermore, the detailed listing of DRMs along with frequencies observed per sample is provided in supplementary file 7.

**Figure 2.**
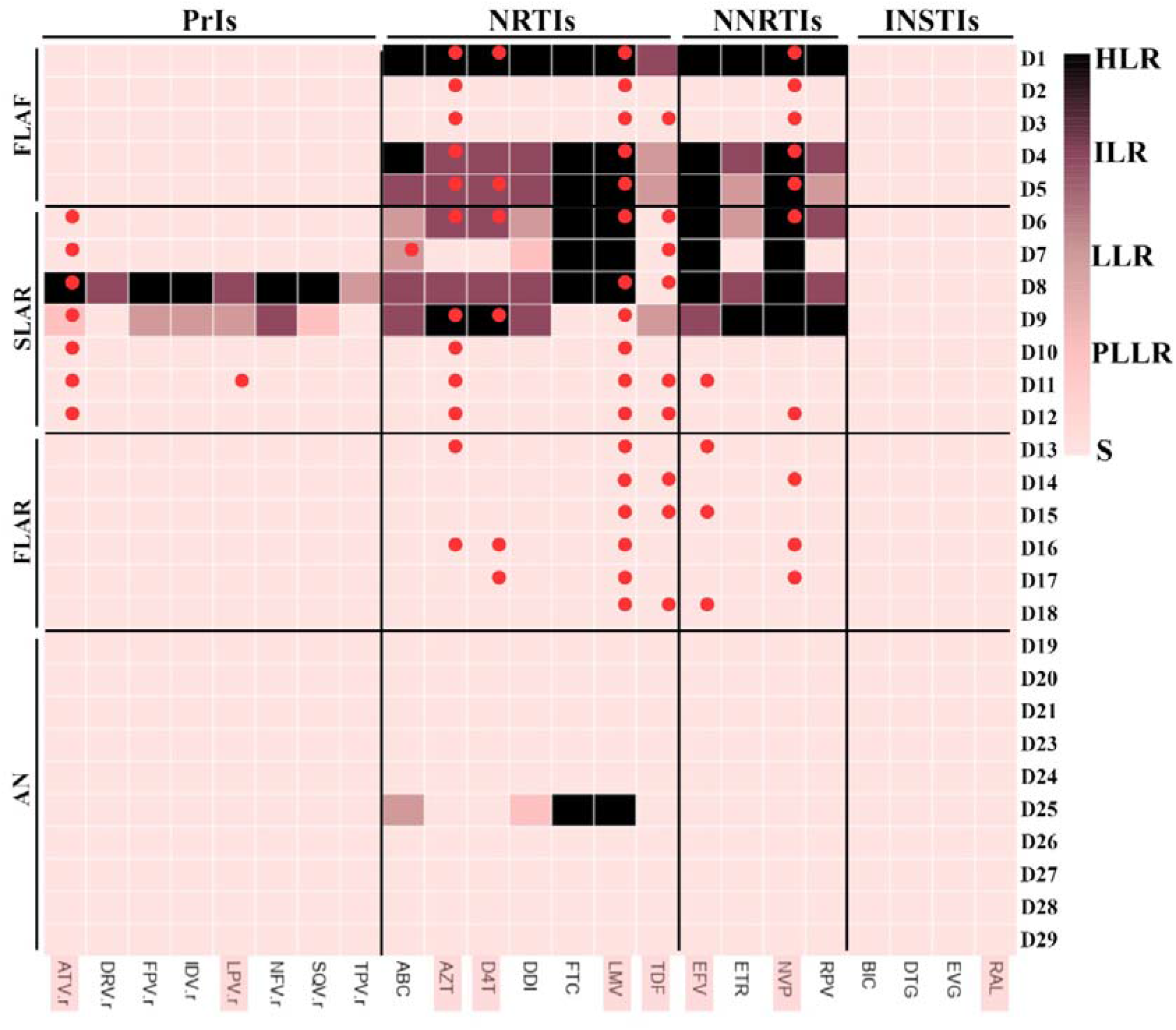
DRM analysis by sanger sequencing: Heatmap of response of individuals S1-S32 (except S25, not tested) to 8 Protease Inhibitors (**PrIs**), 7 Nucleoside reverse transcriptase inhibitors (**NRTIs**), 4 Non-nucleotide reverse transcriptase inhibitors (**NNRTIs**) and 3 Integrase Strand Transfer Inhibitors (**INSTIs**) as a heatmap. Twenty-three antiretroviral drugs have been plotted on the x axis while data from 31 individuals has been plotted on the **Y** axis. Each pixel refers to the predicted response of one individual to one drug colored as per the color-key provided to denote responses ranging from susceptible to high level resistance. Drugs received by the participants as per their clinical records have been indicated by red circles. Furthermore, antiretroviral drugs available under the National AIDS Control Organization, India guidelines have been denoted by orange boxes. **HLR**: High level resistance, **ILR**: Intermediate level resistance, **LLR**: Low level resistance, PLLR: Potential low-level resistance, S: Susceptible.

**Figure 3.**
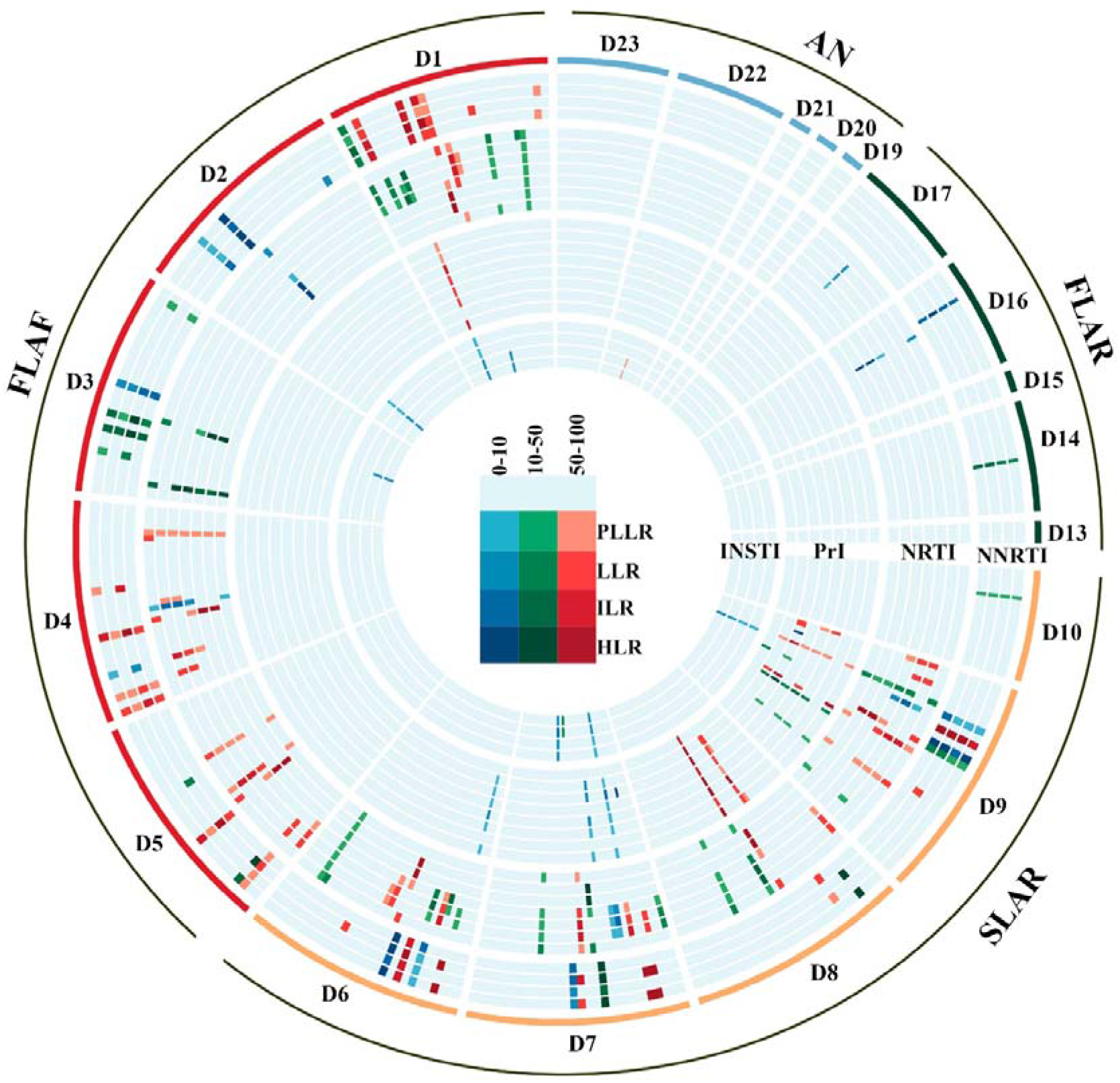
DRM analysis by HTS: Circos heatmap of residue-wise mutation data. The four categories of participants have been indicated by different colored outlines as FLAF, SLAR, FLAR and AN. There are total 23 radially arranged tracks of heatmaps for 23 drugs belonging to the four antiretroviral drug categories viz., in inward order: NNRTI (EFV, ETR, NVP, RPV), NRTI (ABC, AZT, D4T, DDI, FTC, 3TC, TDF), PrI (ATV, DRV, FPV, IDV, LPV, NFV, SQV, TPV) and INSTIs (BIC, DTG, EVG, RAL). The NNRTI tracks indicate 18 individual mutations and 10 mutation combinations, the NRTI tracks show scores for 17 mutations and 16 combinations, PI tracks indicate 24 mutations and 20 combinations whereas INSTI tracks denote results for 21 mutations and 14 combinations.

In the 19 matched datasets, based on presence of specific nucleotide peaks as detected by NovoSNP in electropherograms obtained through SS and SNP detected through HTS, concordance in detection of DRMs was 85.42% for mutations with frequency >50% across sequencing protocols i.e. 41 mutations out of 48 were detected by both technologies. The detection concordance was 48.15% and 3.13% for DRM frequencies between 10-50% and 0-10% respectively. Intra class cross resistance was observed for all the antiviral drug classes by both sequencing methods. Overall, 53 NRTI DRMs were observed in 11 individuals, 10 of which harbored a discriminatory mutation at position 184. Twenty six out of the 53 mutations were TAMs with TAM1 and TAM2 being present in 5 and 8 individuals respectively. In the FLAF group, as expected, Individuals D1, D4 and D5 showed presence of multiple RTI DRMs causing high level resistance along with a few minor mutations (< 50% frequency) causing a low to intermediate level resistance. Individuals D1, D2 and D3 also exhibited presence of DRMs for INSTIs (E138K, G140KR, G163EKR), albeit at a low frequency of <10%. In the SLAR group, all individuals showed presence of DRMs for NNRTIs, NRTIs as well as PrIs. INSTI associated DRMs were also observed in individuals D7 and D9. Individuals from groups FLAR and AN, as expected, only had minor DRMs with a complete absence of any PrI associated DRMs. One high level INSTI DRM (E157Q) was observed in one of the ART naive individual D22. Interestingly, D25, an ART naive individual, exhibited presence of high-level drug resistance mutation towards lamivudine by sanger sequencing suggestive of transmitted drug resistance (Figure 2).

### 3.3. DRM analysis (Quasispecies based)

Prior to performing the quasispecies reconstruction on clinical datasets, a robust benchmarking exercise was undertaken with two experimentally generated control datasets for selection of the optimal algorithm. This was followed by testing an *in-silico* dataset specifically designed to test the ability of the selected reconstruction algorithm to detect DRMs.

#### Benchmarking and Viral Quasispecies reconstruction

Quasispecies reconstruction algorithms (QRAs) ShoRAH, QuRe, PredictHaplo, ViQuaS and QSdpR were selected for the benchmarking analysis [31–36]. To account for error introduced in the sample handling and processing, experimentally generated biological controls were chosen. Benchmarking was performed for a novel control Distributed Concentration Plasmid Mix (DCPM) along with Five virus mix (FVM) dataset (supplementary file 3) [37]. The overall percent identity between viral sequences ranged from 93.54 to 97.46 for FVM while that for DCPM ranged from 90.50 to 95.16. Quasispecies reconstruction was performed for 17 HIV-1 ORFs: p17, p24, p7, p6, Protease, Reverse transcriptase, RNase, Integrase, Vif, Vpr, TatE1, RevE1, Vpu, gp120, gp41, TatE2, and RevE2 by the five selected algorithms for both the controls. Quasispecies data generated was compared with the source viral/plasmid sequences and assessed for recall, precision and processing time.

The overall performance (precision and recall) of all the algorithms was poorer for DCPM with variable predicted frequency of quasispecies (Figure 4). *In silico* mutations generated as artefacts of QRAs can mislead clinical diagnosis, therefore, PredictHaplo with the highest precision (lowest generation of *in silico* recombinants) and high recall for both FVM and DCPM was selected.

**Figure 4.**
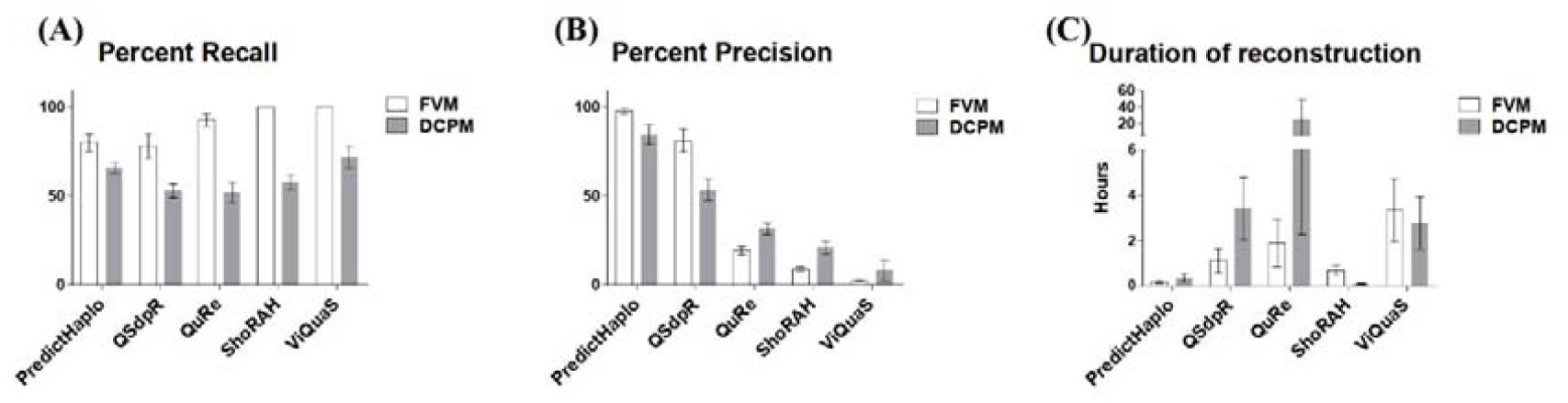
Benchmarking of Viral Quasispecies reconstruction algorithms: Grouped histograms of Benchmarking of 5 Quasispecies reconstruction tools: PredictHaplo, QSdpR, QuRe, ShoRAH and ViQuaS for 17 viral ORFs under 3 categories a) Percent recall b) Percent precision and c) Duration of reconstruction for Five Virus Mix (FVM) and Distributed concentration plasmid mix (DCPM) controls.

Before applying it to the clinical datasets, PredictHaplo was also tested on an *in-silico* NFLG control dataset (ISNC) generated from five variants (frequency: 2-50%, percent identity range: 95-99%) of pIndieC1 genome containing various combinations of DRMs (4 each in PR, RT and IN genes: supplementary file 4). ISNC included a hypermutated sequence with a unique DRM as well as internal stop codons. Consonant with the biological controls, percent recall observed across PR, RT and IN was 60, 80 and 40 respectively (avg: 60%) while percent precision was 100 for all the regions. Furthermore, hypermutated quasispecies, in spite of being present only at 10% abundance, could be correctly reconstructed for all the three genes with detection of not only the unique DRM but also internal stop codons and hypermutations. This confirmed the ability of PredictHaplo to detect DRMs as well as to distinguish those present in hypermutated sequences.

#### Quasispecies Analysis of clinical datasets

Following application to the clinical datasets, reconstructed quasispecies of PR, RT and IN were obtained to be in the range of 1 to 9 per region (supplementary file 8) which were then evaluated for phylogenetic relatedness. All of the quasispecies generated from the same dataset showed monophyletic clustering, demonstrating not only the phylogenetic association of reconstructed quasispecies as expected but also, ruling out any possible cross contamination of the samples during genomic DNA isolation, amplification as well as sequencing. Furthermore, for the three cases of clinically recorded HIV transmission (D16 to D21, D21 to D17 and D14 to D13), phylogenetic association could clearly be established for all 3 genes as indicated in Figure 5 and supplementary file 8. These results demonstrated the ability of PredictHaplo to reconstruct HIV transmission events recorded to have occurred at least 7 years prior to sampling of proviral DNA for the present study.

**Figure 5:**
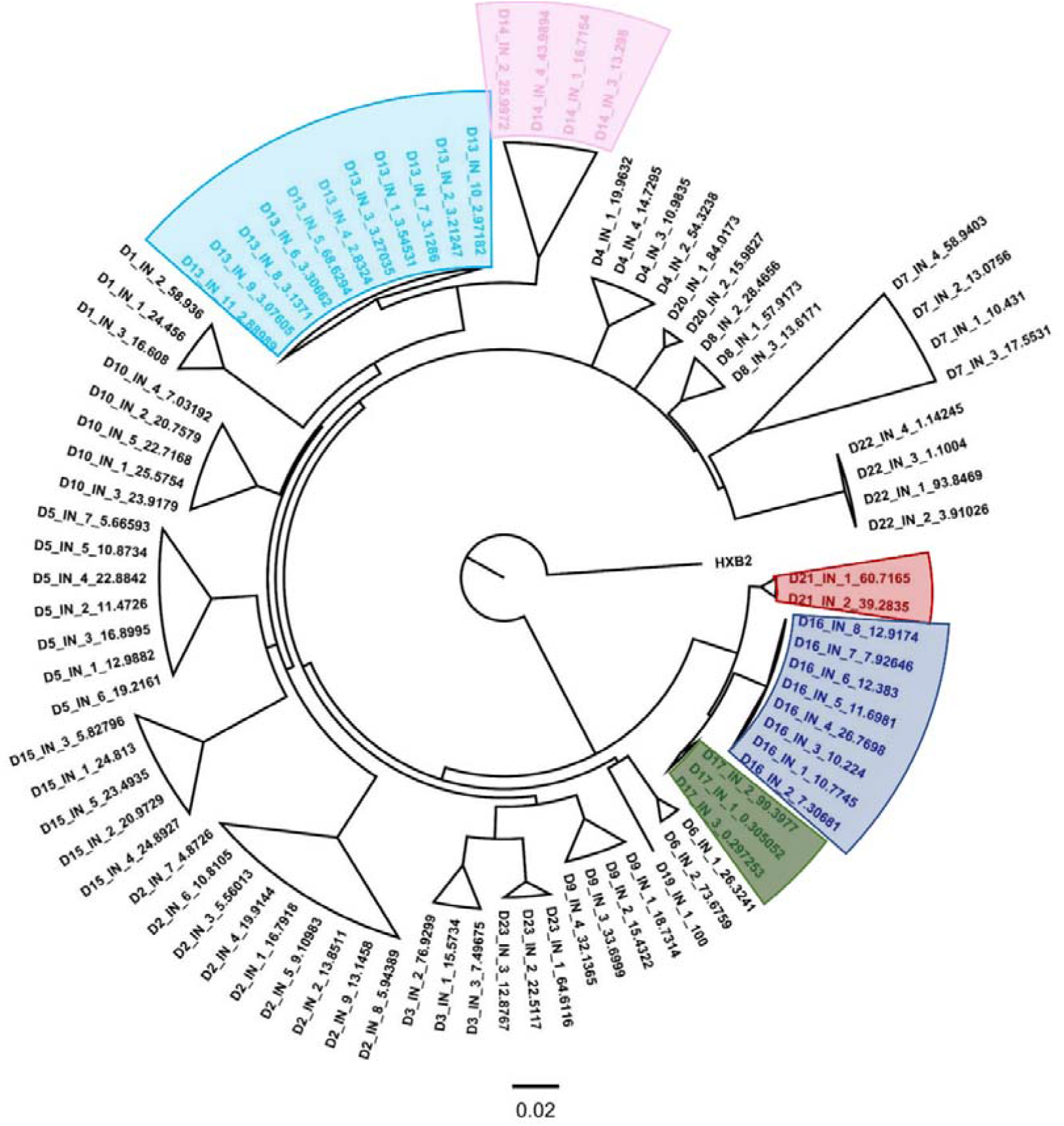
Phylogenetic analysis of quasispecies of integrase gene: Maximum likelihood phylogenetic tree was generated for quasispecies reconstructed for integrase gene from clinical datasets and edited with FigTree. Dataset name codes are in the following format: ‘Dataset-Gene-QuasispeciesNo-Percent Frequency’. Datasets from individuals with shared HIV transmission history have been color coded and highlighted as follows: D16: blue, D17: green. D21: red; D13: sky blue and D14: pink

Prior to DRM analysis, reconstructed quasispecies were assessed for quality/hypermutations/internal stop codons with QC tool of the LANL database. Following rejection of hypermutated sequences, quasispecies were tested with Stanford HIVDb for presence and drug susceptibility of predicted DRMs. RT drug resistance was predicted for 4 out of 5 FLAR individuals and PredictHaplo failed to predict mutations with <10% frequency across categories, represented, for example, in individual D2 (Figure 3 and 6A). Encouragingly however, in contrast to residue wise analysis, this approach was successful in identifying and discarding DRMs on probable hypermutated sequences present at a frequency of ∼20% for individual D7. Concordance evaluation of DRM detection by both technologies (Figure 6B, C and D) showed significant correlation for prediction of PrI DRMs (spearman r=0.92 with p value<0.0001), RTI DRMs (spearman r=0.96 with p value <0.0001) as well as for cumulative DRMs across all samples (spearman r=0.91 with p value<0.0001). In summation, QRA supported DRM analysis was shown to improve accuracy of DRM detection at frequencies >10%.

**Figure 6:**
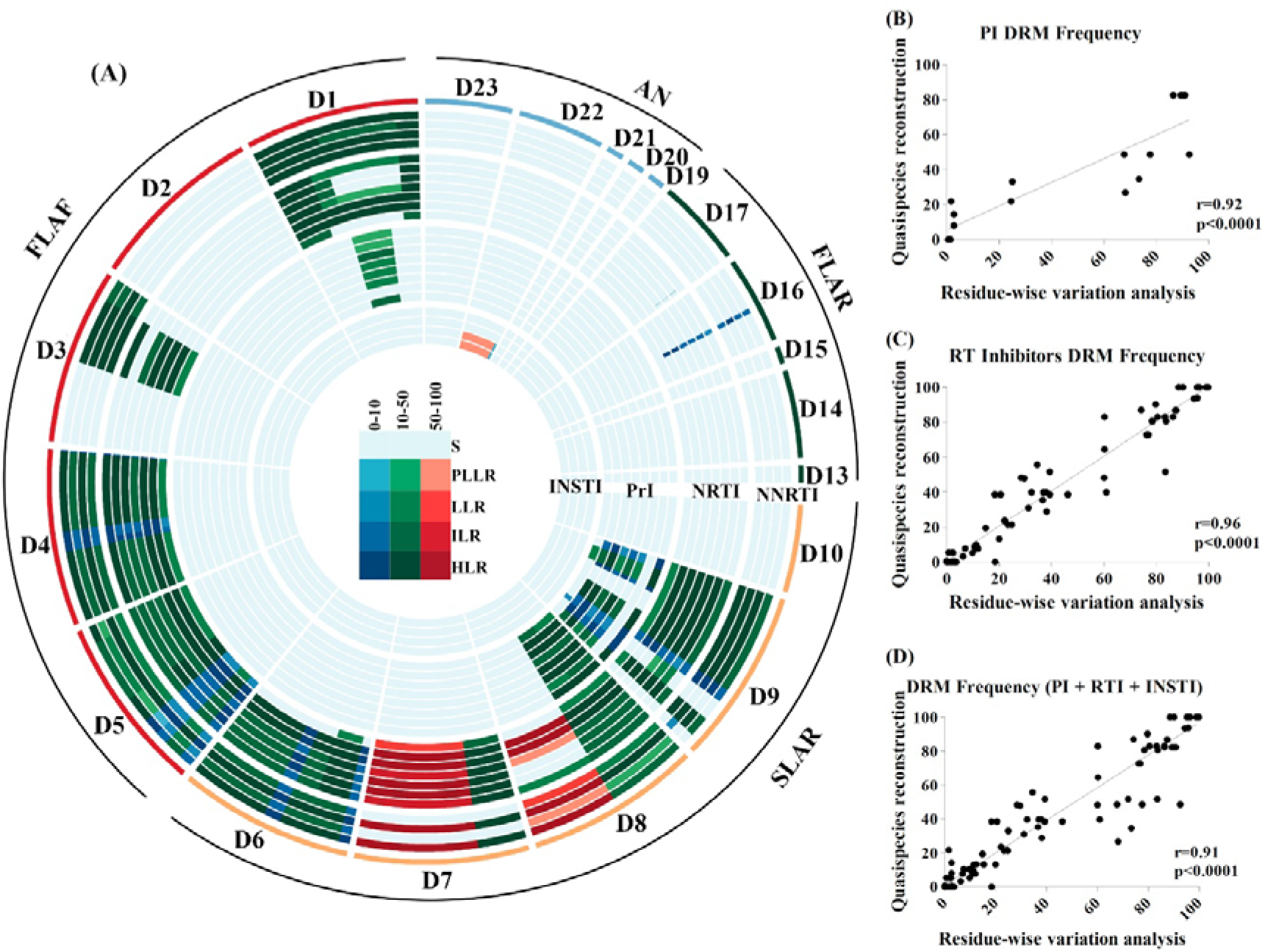
Quasispecies reconstruction-based DRM analysis: A) A heatmap of quasispecies data obtained in a circos format. The four categories of participants have been indicated by different colored outlines as FLAF, SLAR, FLAR and AN. There are total 23 radially arranged tracks of heatmaps for 23 drugs belonging to the four antiretroviral drug categories viz., in inward order: NNRTI (EFV, ETR, NVP, RPV), NRTI (ABC, AZT, D4T, DDI, FTC, 3TC, TDF), PrI (ATV, DRV, FPV, IDV, LPV, NFV, SQV, TPV) and INSTIs (BIC, DTG, EVG, RAL). As per the color key, responses of quasispecies present at >50% predicted frequencies have been denoted in red color spectrum. Similarly, quasispecies with frequency between 10-50% and below 10% are indicated with green and blue color spectra respectively. Instead of different DRMs, this plot indicates overall susceptibility of the sequence, taking into account all possible mutations present in the same viral sequence, therefore, the sequences have been distributed in the drug tracks with their lengths proportional to their predicted frequencies. Correlation analysis between mutation frequency predicted by quasispecies reconstruction (y axis) vs residue wise analysis for B) Protease Inhibitors C) RT inhibitors (NRTI + NNRTI) and D) All DRMs (PrI + NRTI + NNRTI + INSTI)

## 4. Discussion

Success of ‘Treatment as prevention’, depends largely on sensitive DRM monitoring and subsequent adherence support. Suboptimal suppression of viremia due to non-adherence to ART results in emergence of drug resistance. This is a major public health concern especially in resource limited countries such as India, as it may not only result in treatment failure but also lead to transmission of DRMs in a population, limiting future treatment options in therapy naïve individuals [6,49–52]. Present study evaluated the utility of an HTS based and quasispecies reconstruction algorithms (QRA) supported DRM detection approach in HIV-1 clade C infected individuals from Mumbai, India. We believe this is the first report employing open source QRA, to protease, reverse transcriptase and integrase sequences for DRM detection in HIV-1 clade C infected individuals at different stages of antiretroviral therapy mediated viral suppression. Out of the 29 recruited participants, NFLG data was generated through HTS for a subset of 20 participants wherein the entire genome could be successfully amplified and sequenced. To maintain stringency of the HTS workflow, amplification and sequencing artefacts were excluded by using an ‘experimental error control (EEC)’ allowing detection of variant frequencies as low as 0.5%. However, ‘EEC’ was not able to address the bias introduced during isolation of PBMC and subsequent isolation of genomic DNA and may therefore have underreported the threshold. Concordance between DRM detection by sanger and HTS was observed to be proportional to HTS DRM frequency. This disparity at frequencies <50% can be ascribed to lower sensitivity of sanger sequencing as previously reported [14]. HTS based analysis provided an integrated depiction of drug responses, in terms of number of DRM mutations, their frequency and predicted susceptibility to drugs, which is a more clinically relevant interpretation. This representation however, had to be curated manually following careful parsing of the HTS data which limits its applicability in clinical pathology due to complicated implementation. Easily applicable ‘frequency sensitive’ interpretation algorithms need to be developed and thoroughly assessed for analyzing HTS DRM data such as recently published algorithm geno2pheno [ngs-freq] [53]. Regular DRM monitoring of therapy failing individuals would also enable recycling of NRTI drugs for better prognosis of second line regimens [54]. Identical DRMs have been demonstrated to have differential effects across subtypes due distinct naturally occurring polymorphisms [49,55–57]. There also lacks a consensus on effect of minor DRMs on therapy prognoses [58–60]. Therefore, extensive epidemiological surveying of different HIV-1 clades in distinct populations would be indispensable towards development of accurate DRM assessment algorithms.

Interestingly, in the present study, INSTI resistance associated accessory mutations were also detected as minor variants in 5 (25%-E138K, G140KR, G163EKR) and as a major variant in 1 (5%-E157Q) of the assessed individuals, all naive for INSTIs. DRM E157Q is known to be a polymorphic mutation across HIV-1 subtypes [61]. Mutations E138K, G140KR and G163EKR however, are all reported to be nonpolymorphic [62–64]. While G163R from participant D7 (frequency ∼11.5%) could be detected only in hypermutated sequences by PredictHaplo, others could not be similarly distinguished (frequencies <10%) due to lower sensitivity of the reconstruction algorithm. Most of these may also possibly be due to APOBEC3G induced hypermutations and need further examination. These findings are consistent with the recent report of presence of INSTI resistant variants in ART naive HIV-1 clade C sequences in Ethiopia [65], Interestingly, G163KR, a reportedly non polymorphic accessory DRM in subtype C (present in 16 out of 7284 sequences-0.21%) was reported to be at a much higher frequency in both our dataset (3 out of 20) and Indian sequences reported in the LANL HIV sequence database (6 out of 262-2.3%). INSTI drugs have been made available through the national ART program only recently and therefore, G163KR may possibly be a polymorphic mutation for Clade C sequences from India. An expanded sensitive viral surveillance may therefore be necessary to accurately document drug resistance associated mutations in the Indian population.

A growing appreciation of coevolving DRMs and effects of their combinations on viral fitness has made it imperative to examine their epistatic interactions to delineate of DRM pathways and preempt therapy failure [66–68]. Illumina platform of HTS is most popular due to its cost effectiveness and ability to multiplex large number of samples [19,69,70]. However, its applicability to linkage analyses to study epistatic mutations has been limited due to its short-read length [19]. We attempted to assess ability of QRAs to address this caveat through detection of DRM combinations and correlate it to traditional variant calling method specifically for short read HTS data. To select the optimal algorithm for the task and to strengthen the confidence in the predicted sequences, a benchmarking exercise was undertaken. Instead of an *in-silico* control, to account for errors introduced by sample processing, experimentally generated biological controls were selected. Giallonardo *et al* have published and demonstrated applicability of the five-virus control (FVM) for benchmarking such algorithms, wherein five subtype B viral stocks have been mixed in near equimolar concentration, amplified and sequenced [37]. While this is an excellent control for detecting the ability of the tools to decipher mixed viral populations, it does not simulate a clinical sample wherein quasispecies are present also in varying frequencies of minor and major variants. To address this frequency variability, present study developed a new ‘Distributed Concentration Plasmid Mix (DCPM)’ control. This is the first reported HTS dataset of HIV-1 clade C consisting of near full length viral genomes from 5 plasmids mixed in ∼0.5 to 85% frequency range which can be used for benchmarking of all future quasispecies reconstruction algorithms. The algorithm selected in the present study, PredictHaplo, was able to correctly predict DRMs with frequency >10%. Additionally, it could also differentiate DRMs present in hypermutated sequences. PredictHaplo however, was conservative in terms of its recall of viral sequences at frequencies < 10% consistent with earlier reports [37,71]. Therefore, more sensitive algorithms need to be developed to take advantage of increased sensitivity afforded by high throughput sequencing. In spite of the NFLG clinical datasets, Quasispecies analysis reported herein was restricted to generation of distinct variants of PR, RT and IN genes to address effects of intra-protein DRM interactions. While we did try to generate NFLG quasispecies during benchmarking, many of the tools either failed the task or required more computational resources than available. Therefore, the benchmarking and the subsequent analyses was restricted to gene level DRM detection. Consequently, presence of hypermutations and stop codons in other regions of the genome could not be excluded and/or addressed. Also, extension of QR to longer reading frames covering the entire genome could result in detection of novel interprotein correlated mutations and is planned in a future study which would include the present NFLG data.

The present study employed proviral DNA for detection of DRMs, as some of the participants did not have detectable plasma viremia. Addition of matched viral RNA analysis with a separate control dataset to account for RTPCR induced errors would be valuable in further establishing the utility of QRAs in differentiating actively replicating, defective as well as hypermutated viral genomes. This unfortunately could not be addressed in our study due to paucity of plasma samples. While the present study was restricted to HIV-1 clade C, the dominant clade in India, an identical workflow could be applied and tested for other HIV clades. Furthermore, this approach can be extended to other anatomical sites within infected participants to analyze DRMs/mutations influencing compartmentalization and viral fitness of viral quasispecies [72]. Dried blood spots (DBS) are often the only method of specimen collection in rural centers with limited resources and little to no sophisticated amenities to maintain biological samples for long durations. Thus, establishing utility of the workflow described herein for nucleic acids (both RNA and DNA) isolated from DBS would greatly aid in epidemiological monitoring of DRMs in such settings [73,74].

## 5. Conclusions

This study explores a sensitive method to improve existing DRM testing based on quasispecies reconstruction from archived viral sequences. Following further optimization to increase sensitivity, this would be a useful tool for optimal management of ART in PLHIV, especially from resource limited settings.

## Supplementary Materials

The following are available online at www.mdpi.com/xxx/s1, Supplementary file 1: Participant information and therapy duration radial plot, Supplementary file 2: Detailed PCR description, Supplementary file 3: Benchmarking of Quasispecies reconstruction algorithms, Supplementary file 4: Analysis of PredictHaplo with an *in-silico* control dataset, Supplementary file 5: Bash Commands used for analysis of HTS data, Supplementary file 6: Circos heatmap of DRMs from matched sanger datasets, Supplementary file 7: List of DRMs detected from clinical samples, Supplementary file 8: Details of quasispecies reconstructed from clinical samples and their phylogenetic analysis.

### Author Contributions

Conceptualization, Jyoti Sutar and Vainav Patel; Data curation, Jyoti Sutar, Shilpa Bhowmick and Varsha Padwal; Formal analysis, Jyoti Sutar; Funding acquisition, Vainav Patel and Atmaram Bandivdekar; Investigation, Jyoti Sutar, Shilpa Bhowmick and Varsha Padwal; Methodology, Jyoti Sutar; Project administration, Vainav Patel; Resources, Vidya Nagar and Priya Patil; Software, Jyoti Sutar; Supervision, Vainav Patel; Visualization, Jyoti Sutar; Writing – original draft, Jyoti Sutar; Writing – review & editing, Vainav Patel and Atmaram Bandivdekar.

## Acknowledgments

The following reagents were obtained through the NIH AIDS Reagent Program, Division of AIDS, NIAID, NIH: 1) HIV-1 93IN999 Non-Infectious Molecular Clone from Dr. Kavita Lole, Dr. Robert Bollinger, and Dr. Stuart Ray (cat# 3963). 2) HIV-1 94IN476.104 Non-infectious Molecular Clone from Drs. Cynthia M. Rodenburg, Beatrice H. Hahn, Feng Gao, and the Aaron Diamond AIDS Research Center 3) HIV-1 98IN012.14 Infectious Molecular Clone from Drs. Cynthia M. Rodenburg, Beatrice H. Hahn, Feng Gao, and the UNAIDS Network for HIV Isolation and Characterization (cat# 6188). 4) HIV-1 93IN905 Infectious Molecular Clone from Dr. Kavita Lole, Dr. Robert Bollinger, and Dr. Stuart Ray (cat# 3964). The authors would like to acknowledge Dr. Debashis Mitra, NCCS (Pune, India) and Dr. Robin Mukhopadhyay for providing pIndieC1 molecular clone, Mrs. Shilpa Velhal and Ms. Varsha Prabhu for technical assistance in the study. The authors would also like to acknowledge Dr. Somsubhra Barik (University of Texas, USA) for assistance with troubleshooting of QSdpR tool implementation.

---

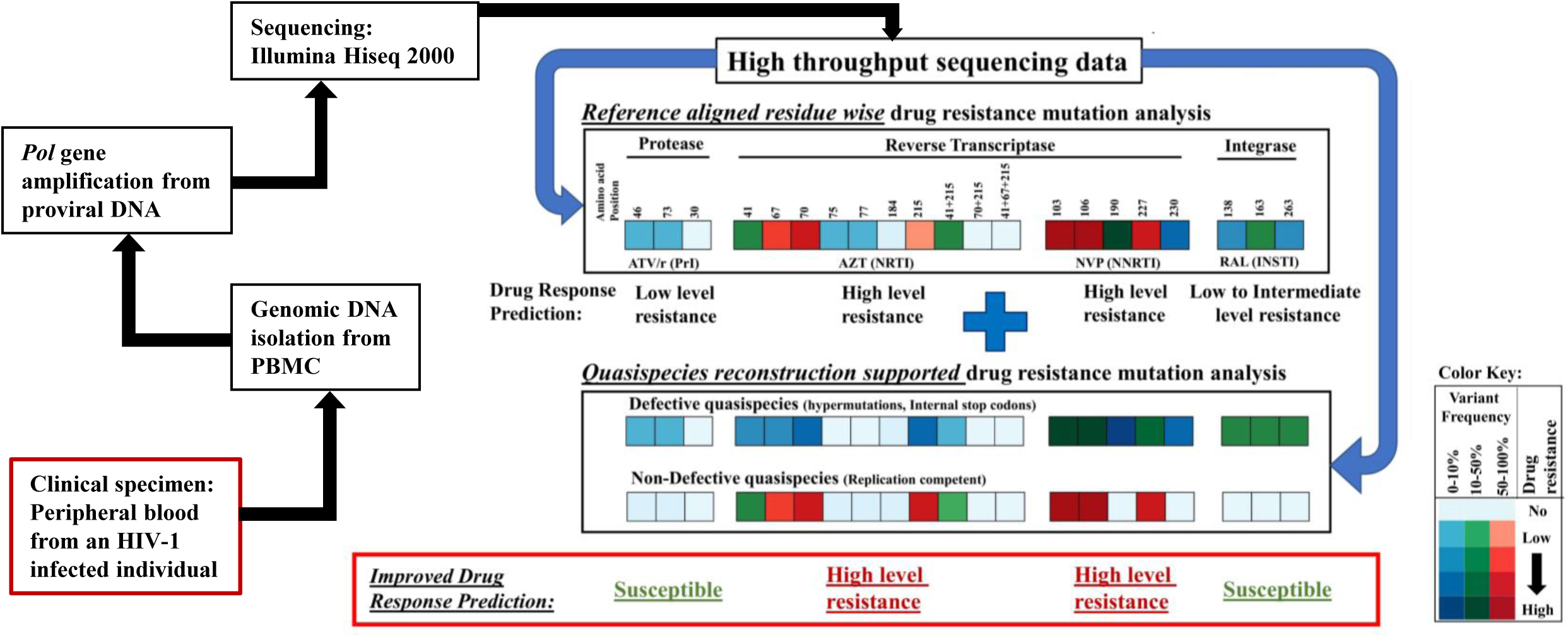

