## Supplementary File 1 for "Improved HIV-1 drug resistance mutation prediction using quasispecies reconstruction supported analysis"

|  | **Age/Sex** | **Last known CD4 Count (cells/mm3)** | **Peak CD4 count (cells/mm3)** | **baseline CD4 count (cells/mm3)** | **Last Viral load (copies/mL)** | **Category** | **Duration of HIV infection (Months)** | **Duration of ART exposure (Months)** | **Current ART combination** | **Antiretroviral drugs received so far** | **Sequenced by** |
| --- | --- | --- | --- | --- | --- | --- | --- | --- | --- | --- | --- |
| D1 | 40/M | 162 | 238 | 135 | 304046 | FLAF | 44 | 26 | ZDV + 3TC + NVP | D4T, ZDV, 3TC, NVP | SS, HTS |
| D2 | 40/M | 372 | 899 | 279 | 184723 | FLAF | 45 | 29 | ZDV + 3TC + NVP | ZDV, 3TC, NVP | SS, HTS |
| D3 | 55/M | 124 | 385 | 86 | 34092 | FLAF | 74 | 74 | TDF + 3TC + NVP | ZDV, 3TC, NVP, TDF | SS, HTS |
| D4 | 48/F | 269 | 1022 | 386 | 7293 | FLAF | 168 | 108 | ZDV + 3TC + NVP | ZDV, 3TC, NVP | SS, HTS |
| D5 | 36/F | 179 | 547 | 155 | 29194 | FLAF | 228 | 108 | D4T + 3TC + NVP | ZDV, 3TC, NVP | SS, HTS |
| D6 | 48/M | 414 | 879 | 195 | <34 | SLAR | 108 | 108 | TDF + 3TC + ATV/r | D4T, ZDV, 3TC, NVP, TDF, ATV/r | SS, HTS |
| D7 | 51/M | 194 | 353 | 17 | 97 | SLAR | 99 | 75 | TDF + ATV/r + ABC | TDF, ATV/r, ABC | SS, HTS |
| D8 | 45/M | 185 | 185 | NAV | 848 | SLAR | 76 | 40 | TDF + 3TC + ATV/r | TDF, 3TC, ATV/r | SS, HTS |
| D9 | 45/F | 202 | 202 | 38 | 132788 | SLAR | 200 | 60 | D4T + 3TC + ATV/r | D4T, 3TC, ATV/r | SS, HTS |
| D10 | 42/F | 1684 | 1684 | 827 | TND | SLAR | 26 | 24 | ZDV + 3TC + ATV/r | ZDV, 3TC, ATV/r | SS, HTS |
| D11 | 59/M | 194 | 512 | 164 | TND | SLAR | 90 | 84 | TDF + 3TC + ATV/r | ZDV, 3TC, EFV, TDF, ATV/r, LPV/r | SS |
| D12 | 32/M | 118 | 118 | 107 | <34 | SLAR | 125 | 120 | TDF + 3TC + ATV/r | ZDV, NVP, TDF, 3TC, ATV/r | SS |
| D13 | 37/F | 1061 | 1061 | 679 | TND | FLAR | 180 | 68 | ZDV + 3TC + EFV | ZDV, 3TC, EFV | SS, HTS |
| D14 | 43/M | 714 | 746 | 590 | <34 | FLAR | 180 | 94 | TDF + 3TC + NVP | TDF, 3TC, NVP | SS, HTS |
| D15 | 45/F | 424 | 597 | 73 | <34 | FLAR | 21 | 13 | TDF + 3TC + EFV | TDF, 3TC, EFV | SS, HTS |
| D16 | 45/M | 864 | 898 | 437 | 457 | FLAR | 76 | 66 | ZDV + 3TC + NVP | D4T, ZDV, 3TC, NVP | SS, HTS |
| D17 | 9/M | 1324 | 1324 | NA | TND | FLAR | 70 | 63 | D4T + 3TC + NVP | D4T, 3TC, NVP | SS, HTS |
| D18 | 35/F | 444 | 444 | NA | TND | FLAR | 24 | 12 | TDF + 3TC + EFV | TDF, 3TC, EFV | SS |
| D19 | 28/M | 534 | NA | NA | 197054 | AN | 6 | NA | NA | NA | SS, HTS |
| D20 | 45/M | 612 | NA | NA | 6585 | AN | 30 | NA | NA | NA | SS, HTS |
| D21 | 40/F | 440 | NA | NA | 2749 | AN | 48 | NA | NA | NA | SS, HTS |
| D22 | 43/M | 527 | NA | NA | 861484 | AN | 60 | NA | NA | NA | HTS |
| D23 | 35/M | 230 | NA | NA | 101032 | AN | 4 | NA | NA | NA | SS, HTS |
| D24 | 25/F | 510 | NA | NA | 1370 | AN | 48 | NA | NA | NA | SS |
| D25 | 36/M | 418 | NA | NA | 32851 | AN | 24 | NA | NA | NA | SS |
| D26 | 34/F | 548 | NA | NA | 3456 | AN | 36 | NA | NA | NA | SS |
| D27 | 37/F | 536 | NA | NA | TND | AN | 28 | NA | NA | NA | SS |
| D28 | 53/F | 762 | NA | NA | 32694 | AN | 48 | NA | NA | NA | SS |
| D29 | 34/F | 715 | NA | NA | 668998 | AN | 36 | NA | NA | NA | SS |

**Footnote:** FLAF: First line ART failing, SLAR: Second line ART receiving, FLAR: First line ART receiving, AN: ART naive, NAV: Not available, TND: Target not detected, NA: Not applicable, ZDV: Zidovudine, D4T: Stavudine, ABC: Abacavir, 3TC: Lamivudine, NVP: Nevirapine, EFV: Efavirenz, TDF: Tenofovir disoproxil fumarate, Ritonavir boosted Atazanavir: ATV/r, Ritonavir boosted lopinavir: LPV/r. SS: Sanger Sequencing, HTS: High throughput sequencing

**Detailed Therapy duration:**

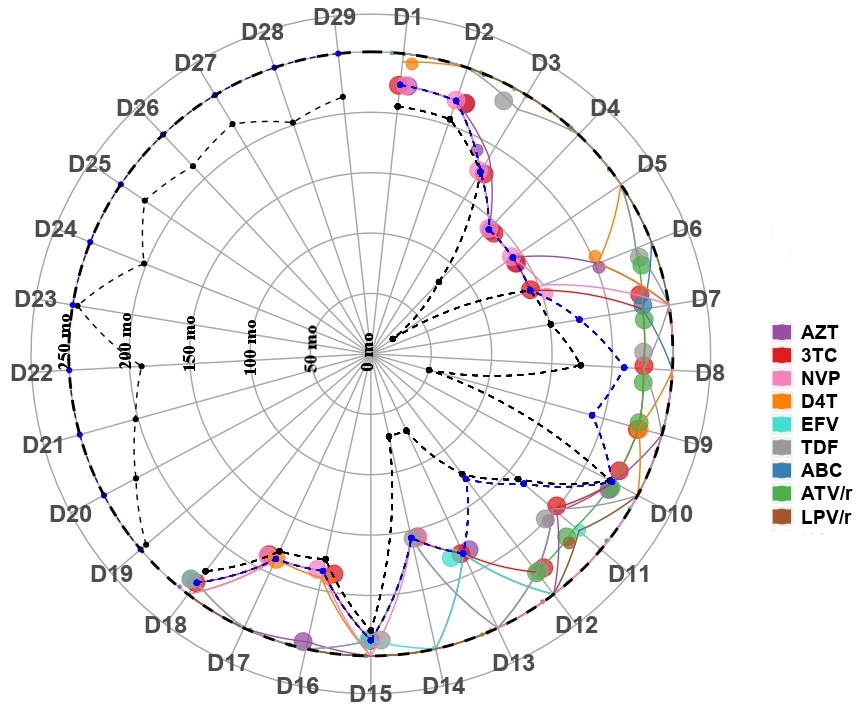

**Therapy Duration:** Radial plot indicating detailed therapy duration for each participant arranged radially. Y axis (drawn outward) indicates period (months). All the drug points have been color coded as indicated in the color key. Most recent therapy is indicated with largest circles decreasing in order of past regimens. Black dashed line and blue dashed line indicate period since detection and therapy initiation respectively.
