## Supplementary File 2 for "Improved HIV-1 drug resistance mutation prediction using quasispecies reconstruction supported analysis"

**Detailed PCR description**

**Protease (PR), Reverse Transcriptase (RT) and Integrase (IN) genes:**

PR (HXB2: 2147-2639), RT (HXB2: 2511-3351) and IN (HXB2: 4343-5092) regions were amplified with a nested PCR approach with a common first-round ‘pol amplification’ protocol (Nadai et al., 2008) using Primestar polymerase (TaKaRa). Subsequently, PR, RT and IN were amplified with 5µL of the 1st round product with primers indicated in Table S1 with GeneAll polymerase enzyme (GeneAll, Korea) as described in table S2.

**Near full length genome (NFLG) PCR amplification:**

NFLG was amplified as reported by Nadai et al(Nadai et al., 2008) with 3 overlapping fragments covering gag, pol and env regions of HIV-1 genome (HXB2: 769-9089). using Primestar polymerase (TaKaRa) as described in Table S1 with primer sets described in table S2.

**Table S1**

| **Reagents** | **1^st^ Round Pol PCR as well as NFLG:3 amplicon strategy (1X)**  **PrimeStar polymersase (TaKaRa)** | **Protease, Reverse Transcriptase and Integrase 2^nd^ round PCR (1X)**  **GeneAll polymerase enzyme (GeneAll)** |
| --- | --- | --- |
| **PCR buffer** | 1X | 1X |
| **DNTP mix** | 1.6mM | 1mM |
| **MgCl2** | 2.75mM (present in the buffer) | 3.6mM |
| **Forward and reverse primers** | 0.4µM each | 0.4µM each |
| **Polymerase enzyme** | 5U | 5U |
| **Template** | ~1µg of DNA/5µL of first round product | 5µL of first round product |
| **Nuclease Free water** | To make up the volume | To make up the volume |
| **Cycling Conditions** | 94^0^ C for 2 mins,  10 cycles of: 94^0^C 10 s, 60^0^C 30 s, 68^0^C 4 mins,  20 cycles of: 94^0^C 10 s, 55^0^C 30 s, 68^0^C 4 mins,  68^0^C for 10 mins | 95^0^C for 5 mins,  40 cycles of: 95^0^C 45 s, 60^0^C 45 s, 72^0^C 1 min,  72^0^C for 10 mins. |

**Table S2**

| **Amplicon** | **Name** | **Sequence (5’ -3’)** | **HXB2 coordinates** | **Direction** | **Amplicon Size (bp)** |
| --- | --- | --- | --- | --- | --- |
| 1^st^ round Pol/ 1^st^ round pol fragment (3AS) | PolF1 | AGAAATTGCAGGGCCCCTAGGAA | 1996-2018 | Forward | 4357 |
|  | PolR1 | GGTACCCCATAATAGACTGTRACCCACAA | 6324-6352 | Reverse |  |
| Protease | ProteaseF | ACCAGAGCCAACAGCCCCACCA | 2147-2168 | Forward | 493 |
|  | ProteaseR | CCTTTGGGCCATCCATTCCTGGC | 2616-2639 | Reverse |  |
| Reverse Transcriptase | RTF | AGGACCTACRCCTGTCAACATAATTGG | 2511-2538 | Forward | 841 |
|  | RTR | TGTATGTCATTGACAGTCCAGCTG | 3327-3351 | Reverse |  |
| Integrase | INF | TAGCTGTGATCAATGTCAGTTAAAA | 4343-4367 | Forward | 750 |
|  | INR | TCTTCATCCTGTCTACCTGCC | 4072-5092 | Reverse |  |
| 2^nd^ round pol fragment (3AS) | PolF2 | AGANCAGAGCCAACAGCCCCACCA | 2143-2166 | Forward | 4089 |
|  | PolR2 | CTCTCATTGCCACTGTCTTCTGCTC | 6207-6231 | Reverse |  |
| 1^st^ round gag fragment (3AS) | gagF1 | AAATCTCTAGCAGTGGCGCCCGAACAG | 623-649 | Forward | 2855 |
|  | gagR1 | GAATCTCTCTGTTTTCTGCCAGTTC | 3453-3477 | Reverse |  |
| 2^nd^ round gag fragment (3AS) | gagF2 | GCGGAGGCTAGAAGGAGAGAGATGG | 769-793 | Forward | 2570 |
|  | gagR2 | TTTCCCCACTAACTTCTGTATGTCATTGACA | 3308-3338 | Reverse |  |
| 1^st^ round env fragment (3AS) | envF1 | AGARGAYAGATGGAACAAGCCCCAG | 5550-5574 | Forward | 3632 |
|  | envR1 | GTGTGTAGTTCTGCCAATCAGGGAA | 9157-9181 | Reverse |  |
| 2^nd^ round env fragment (3AS) | envF2 | TTAGGCATCTCCTATGGCAGGAAGAAGCGG | 5957-5986 | Forward | 3133 |
|  | envR2 | TCCAGTCCCCCCTTTTCTTTTAAAAA | 9064-9089 | Reverse |  |

Footnote: 3AS: 3 amplicon strategy
