## Supplementary File 3 for "Improved HIV-1 drug resistance mutation prediction using quasispecies reconstruction supported analysis"

**Benchmarking of Quasispecies reconstruction algorithms**

The data presented here describe a benchmarking study to find the most optimal algorithm for application to the clinical data. Five Quasispecies reconstruction tools: ShoRAH, QuRe, PredictHaplo, ViQuaS and more recently published QSdPR were selected for the benchmarking analysis(Barik et al., 2018; Jayasundara et al., 2015; McElroy et al., 2013; Prabhakaran et al., 2014; Prosperi and Salemi, 2012; Zagordi et al., 2011). Instead of an *in silico* control, to account for errors introduced by sample handling as well as processing, experimentally generated biological controls were selected. Giallonardo *et al* have published and demonstrated applicability of the five-virus mix control (FVM) for benchmarking such algorithms, wherein five subtype B viral stocks (HXB2, JR-CSF, NL4-3, YU2 and 89,6) have been mixed in near equimolar concentrations, amplified and sequenced(Di Giallonardo et al., 2014). While this is an excellent control for detecting the ability of the tools to decipher mixed viral populations, it does not imitate a clinical sample wherein quasispecies are present not only in equal but also varying frequencies of minor and major variants. To address the frequency variability, a new control: Distributed Concentration Plasmid Mix (DCPM) was developed in the present study. Quasispecies reconstruction was performed for 17 HIV-1 ORFs: p17, p24, p7, p6, Protease, Reverse transcriptase, RNase, Integrase, Vif, Vpr, TatE1, RevE1, Vpu, gp120, gp41, TatE2, and RevE2 by the five selected algorithms for both the controls.

**Percent Identity**

Percent identity metrices were generated using clustal omega webtool(Sievers et al., 2011). The overall percent identity between viral sequences ranged from 93.54 to 97.46 for FVM while that for DCPM ranged from 90.50 to 95.16 (tables 1A and 1B)

Table 1A

|  | 89.6 | JRCSF | YU2 | HXB2 | NL4-3 |
| --- | --- | --- | --- | --- | --- |
| 89.6 | 100 | 93.81 | 93.60 | 93.89 | 93.54 |
| JRCSF | 93.81 | 100 | 94.63 | 94.57 | 94.39 |
| YU2 | 93.60 | 94.63 | 100 | 95.16 | 94.83 |
| HXB2 | 93.89 | 94.57 | 95.16 | 100 | 97.46 |
| NL4-3 | 93.54 | 94.39 | 94.83 | 97.46 | 100 |

Table 1B

|  | pIndieC1 | p93IN999 | p98IN012.14 | p94IN476.104 | p93IN905 |
| --- | --- | --- | --- | --- | --- |
| pIndieC1 | 100 | 91.35 | 90.50 | 90.11 | 92.42 |
| p93IN999 | 91.35 | 100 | 94.39 | 92.45 | 95.12 |
| p98IN012.14 | 90.50 | 94.39 | 100 | 92.32 | 94.03 |
| p94IN476.104 | 90.11 | 92.45 | 92.32 | 100 | 92.69 |
| p93IN905 | 92.42 | 95.12 | 94.03 | 92.69 | 100 |

**Benchmarking**

**Number of Quasispecies reconstructed**

Expected number of quasispecies generated for both the controls were 5. Number of quasispecies generated were consistently higher for FVM than DCPM except for QSdpR (Figure 1). Overall, for the 17 viral ORFs (expected total: DCPM:85, FVM:85), ViQuaS generated most number of quasispecies (Median values (range):- DCPM:142 (1-635), FVM:221(107-705), Total: DCPM:3474, FVM:705) while PredictHaplo (Median values (range):- DCPM:4(3-9), FVM:4(1-5), Total: DCPM:73, FVM:70) and QSdpR (Median values (range):- DCPM:6(2-8), FVM:5(1-6), Total: DCPM:95, FVM:83) generated the least number of quasispecies. QuRe was unable to reconstruct any quasispecies for gp120 ORF (1513bp) for DCPM.

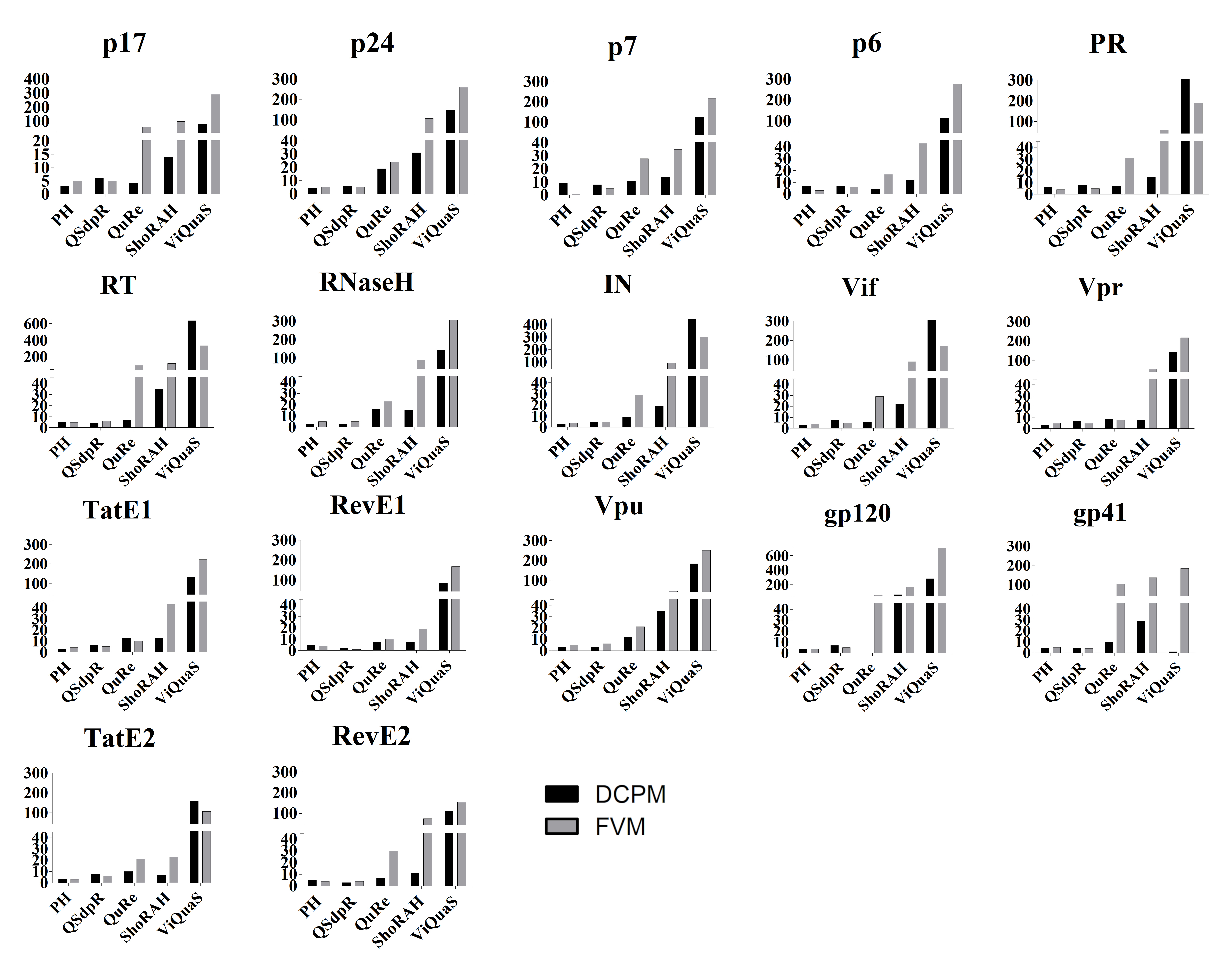
**Figure 1**

**Number of quasispecies generated:** No. of quasispecies have been plotted on the Y axis while the tools have been plotted on the x axis. The bars have been color coded as depicted in the legend

**Benchmarking of Viral Quasispecies reconstruction algorithms*:***

Quasispecies data generated was compared with the source viral/plasmid sequences and assessed for recall, precision and processing time as follows:

1) Recall: (No. of Reconstructed quasispecies closest to the source sequences i.e. accepted quasispecies /expected no of quasispecies) x 100

2) Precision: (No of accepted quasispecies/Total no. of quasispecies generated by the tool) x 100

3) Processing Time: Total duration of quasispecies reconstruction of Protease, Reverse transcriptase and Integrase genes on an intel i7 4790 @ 3.60 GHz (8 threads) processor with 20GB RAM

**Percent Recall**

Expected percent recall for both the controls was 100. Overall, as depicted in figure 2, percent recall for FVM was higher (Median: 100, range: 20-100) than DCPM (Median: 60, range: 0-100). ViQuaS had the highest recall for both the controls (Median values (range), DCPM: 80(20-100), FVM: 100(100-100)) while PredictHaplo had the lowest recall for both the controls (Median values (range), DCPM: 60(60-100), FVM: 80(20-100)).

**Figure 2**

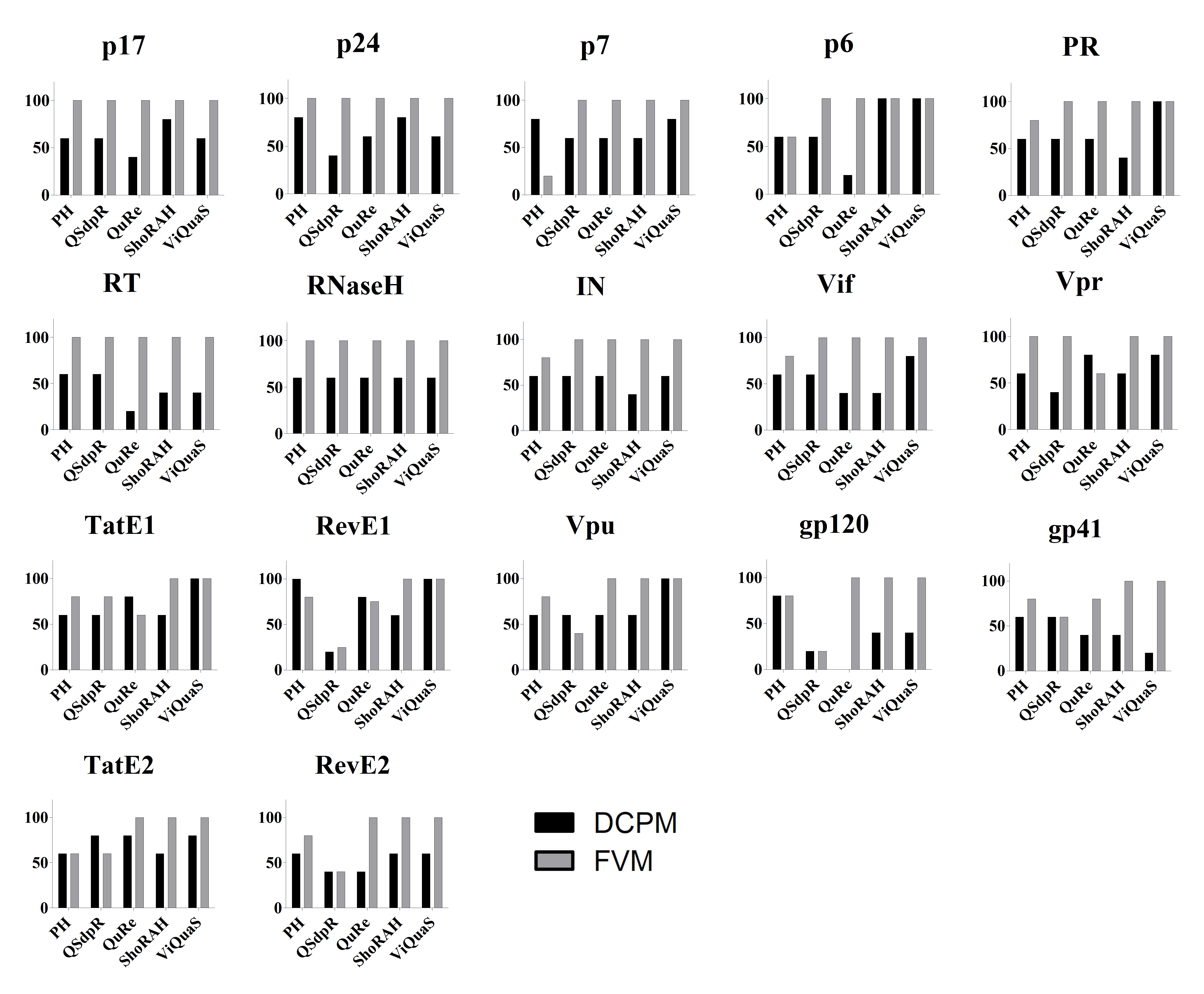
Percent Recall: Percent recall has been plotted on y axis while the tools have been depicted on the X axis. The bars have been color coded as depicted in the legend.

**Percent Precision**

The expected percent precision for both controls was 100. As depicted in figure 3, PredictHaplo was observed to be most precise for both of the controls (Median (range):- DCPM: 100(42.85-100), FVM: 100(80-100)) while ViQuaS was least precise (Median (range):- DCPM:2.67(0.314-100), FVM:2.26(0.70-4.67)). QSdpR had high precision for FVM control (Median: 100, range: 20-100) while that for DCPM was lower (Median: 50, range: 14.28-100). This trend was observed to be consistent for tools QuRe and ShoRAH.

**Figure 3**

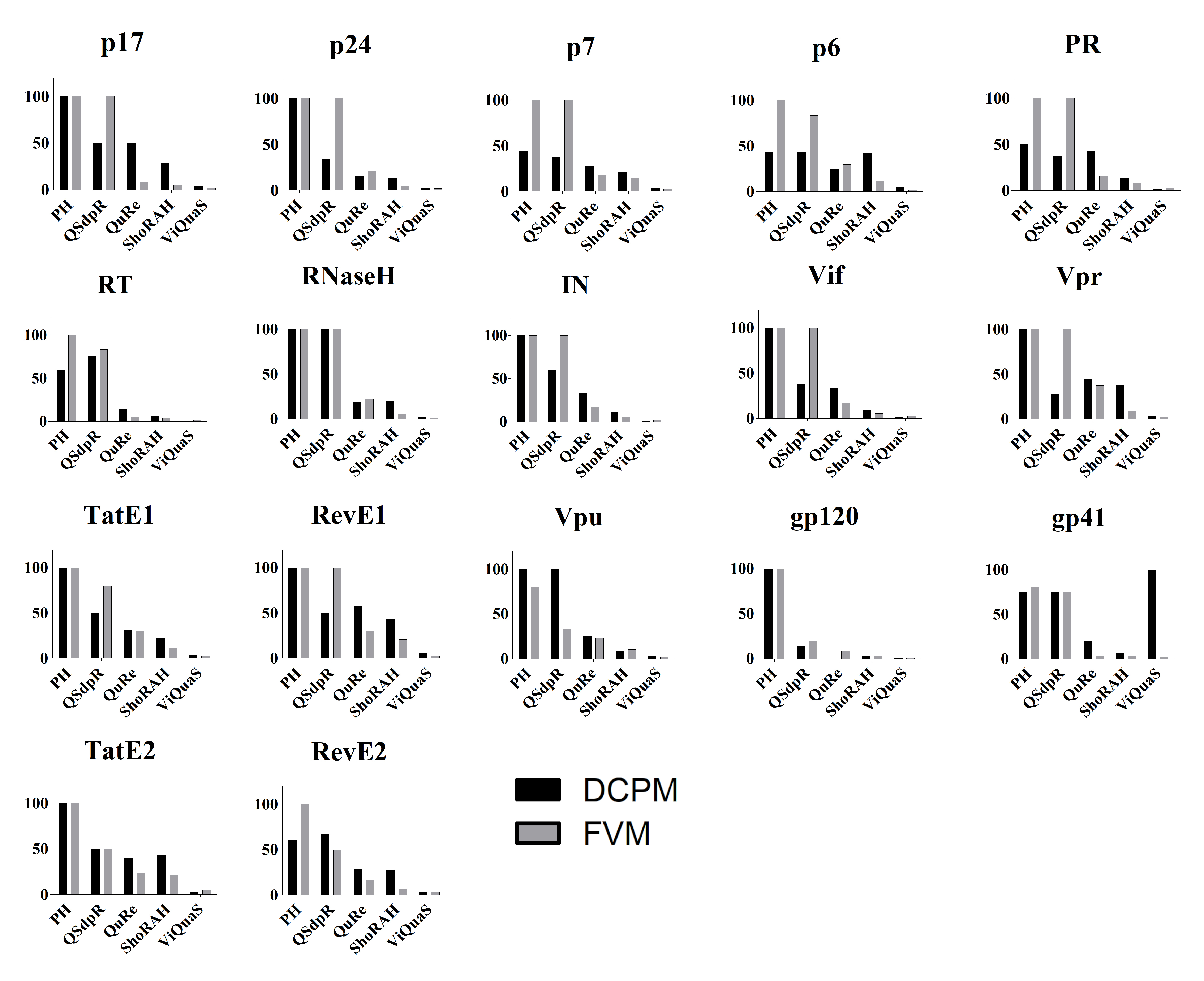
**Percent Precision:** Percent precision has been plotted on y axis while the tools have been depicted on the X axis. The bars have been color coded as depicted in the legend.

**Duration of quasispecies reconstruction**

Quasispecies reconstruction tools with least computational resource requirements i.e. the quickest to reconstruct quasispecies on a given system are expected to be the most efficient and accessible. As depicted in Figure 4, this parameter was assessed (in hours) for PR, RT and IN ORFs only. PredictHaplo reconstructed quasispecies the quickest for both the controls (Median (range):- DCPM:0.24(0.08-0.73), FVM:0.23(0.04-0.25)) while ViQuaS was observed to be the slowest (Median (range):- DCPM:1.9(1.35-5.12), FVM:4.57(0.58-4.89)).

**Figure 4**

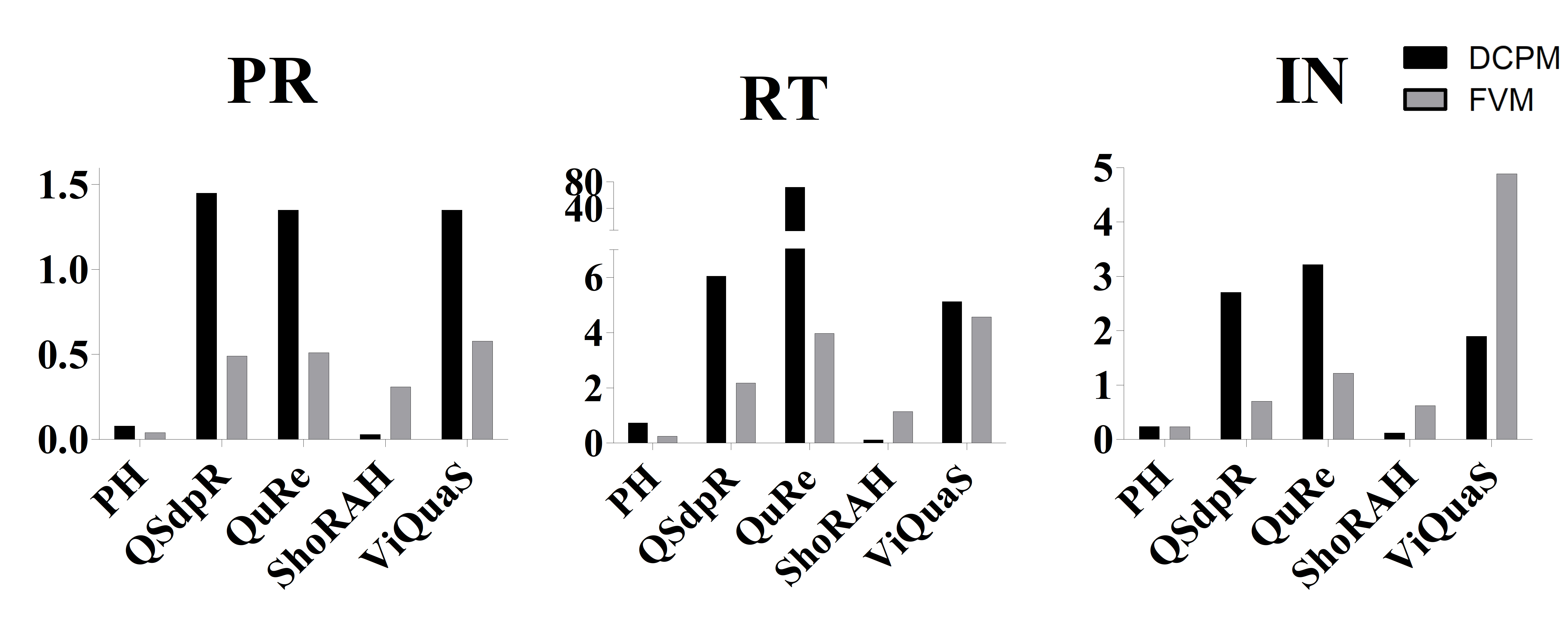

**Duration of quasispecies reconstruction:** Duration (time in hours) has been plotted on y axis while the tools have been depicted on the X axis. The bars have been color coded as depicted in the legend.
