## Supplementary File 4 for "Improved HIV-1 drug resistance mutation prediction using quasispecies reconstruction supported analysis"

**Analysis of PredictHaplo with an *in-silico* control (INSC) dataset:**

**INSC composition:**

| **Sequence Name** | **Abundance (%)** | **PR Mutations** | **RT Mutations** | **IN Mutations** |
| --- | --- | --- | --- | --- |
| pIndieC1_NM (NM) | 30 | None | None | None |
| pIndieC1_hym (hym) | 10 | G48K | L210W | G140S |
| pIndieC1_1M (1M) | 50 | M46I | M184V | E138K |
| pIndieC1_2M (2M) | 8 | D30N, M46I | D67N, M184V | E138K, G163R |
| pIndieC1_3M (3M) | 2 | D30N, M46I, L90M | D67N, Y181C, M184V | E138K, G163R, R263K |

Other than the drug resistance mutations, random synonymous mutations were added in the pIndieC1 sequence to generate up to 5% dissimilarity between the sequences.

In the **pIndieC1_hym sequence**, following hypermutations were added:

**PR:** D25N, G40R, G48K,

**RT:** G152K, G213K, W406* (stop codon)

**IN:** W19*(stop codon), G47K, G59E, G70K, G106K, W132* (stop codon)

Final gene-wise percent identity matrix generated with clustal omega.

|  | **PR** | | | | | **RT** | | | | | **IN** | | | | |
| --- | --- | --- | --- | --- | --- | --- | --- | --- | --- | --- | --- | --- | --- | --- | --- |
|  | hym | 3M | 1M | NM | 2M | hym | 3M | 1M | NM | 2M | hym | 3M | 1M | NM | 2M |
| hym | 100 | 94.95 | 94.95 | 97.31 | 95.29 | 100 | 97.12 | 97.20 | 98.48 | 97.05 | 100 | 96.54 | 96.77 | 98.15 | 96.66 |
| 3M | 94.95 | 100 | 95.96 | 97.64 | 96.97 | 97.12 | 100 | 97.50 | 98.64 | 97.58 | 96.54 | 100 | 97.23 | 98.39 | 97.35 |
| 1M | 94.95 | 95.96 | 100 | 97.64 | 96.63 | 97.20 | 97.50 | 100 | 98.71 | 97.42 | 96.77 | 97.23 | 100 | 98.62 | 97.35 |
| NM | 97.31 | 97.64 | 97.64 | 100 | 97.98 | 98.48 | 98.64 | 98.71 | 100 | 98.56 | 98.15 | 98.39 | 98.62 | 100 | 98.50 |
| 2M | 95.29 | 96.97 | 96.63 | 97.98 | 100 | 97.05 | 97.58 | 97.42 | 98.56 | 100 | 96.66 | 97.35 | 97.35 | 98.50 | 100 |

**Quasispecies reconstruction results:**

| **Gene** | **Expected no. of quasispecies**  **(E)** | **No. of quasispecies generated (G)** | **Correctly predicted quasispecies based on DRMs (C)** | **Percent Recall**  **[formula: (C/E)*100]** | **Percent precision**  **[formula: (C/G)*100]** |
| --- | --- | --- | --- | --- | --- |
| PR | 5 | 3 | 3  [NM, 1M, hym] | 60 | 100 |
| RT | 5 | 4 | 4  [NM, 1M, hym, 2M] | 80 | 100 |
| IN | 5 | 2 | 2  [1M, hym] | 40 | 100 |

**Features detected in the reconstructed hypermutated sequences:**

**PR:** D25N, G40R, G48K.

**RT:** G152K, G213K

**IN:** W19*(stop codon), G47K, G59E, G70K, G106K, W132*(stop codon)
