## Supplementary File 6 for "Improved HIV-1 drug resistance mutation prediction using quasispecies reconstruction supported analysis"

**Circos heatmap of DRMs from matched sanger datasets**

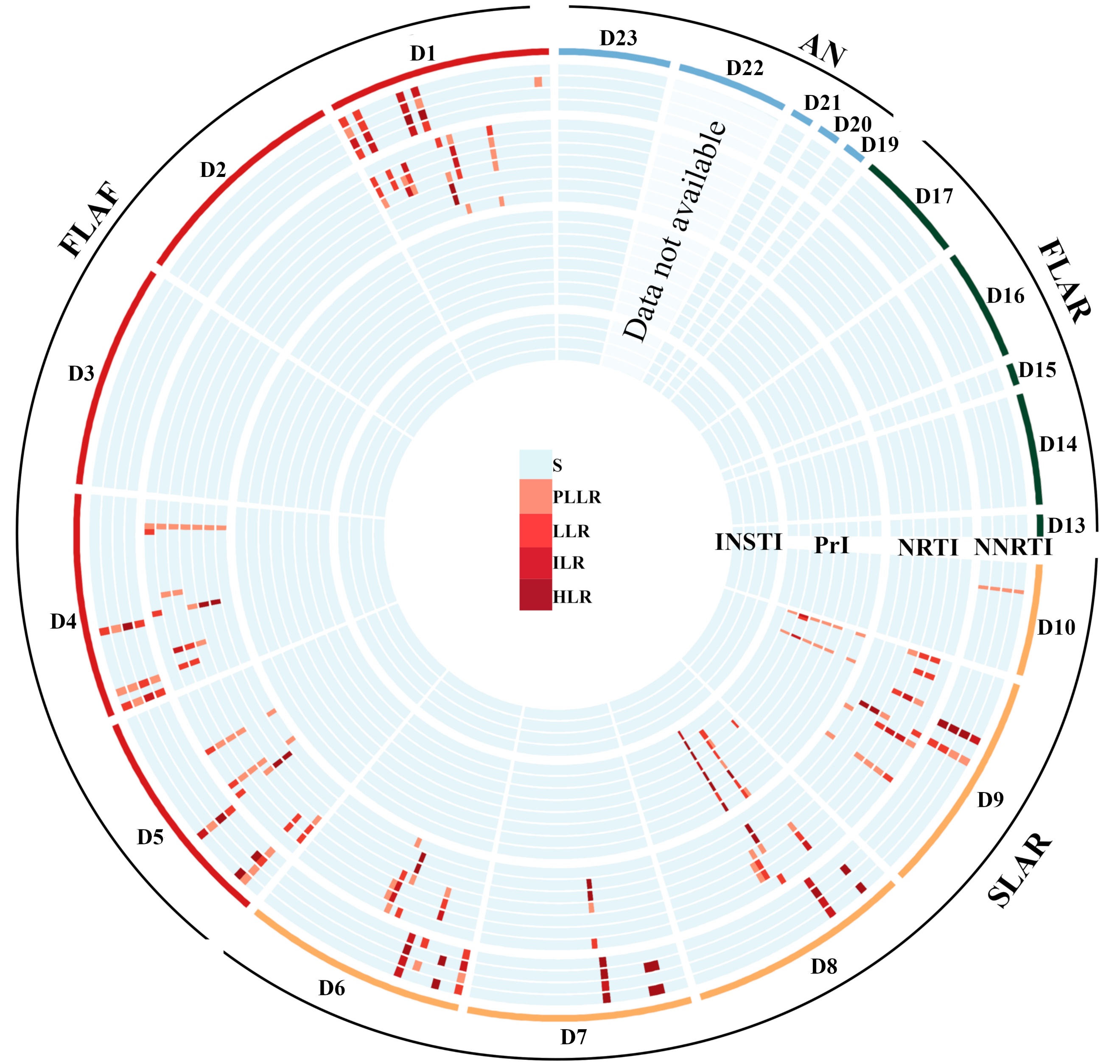

**DRM analysis by Sanger Sequencing:** Circos heatmap of residue-wise mutation data. The four categories of participants have been indicated by different colored outlines as FLAF, SLAR, FLAR and AN. There are total 23 radially arranged tracks of heatmaps for 23 drugs belonging to the four antiretroviral drug categories viz., in inward order: NNRTI (EFV, ETR, NVP, RPV), NRTI (ABC, AZT, D4T, DDI, FTC, 3TC, TDF), PrI (ATV, DRV, FPV, IDV, LPV, NFV, SQV, TPV) and INSTIs (BIC, DTG, EVG, RAL). The NNRTI tracks indicate 18 individual mutations and 10 mutation combinations, the NRTI tracks show scores for 17 mutations and 16 combinations, PI tracks indicate 24 mutations and 20 combinations whereas INSTI tracks denote results for 21 mutations and 14 combinations.
