## Supplementary File 8 for "Improved HIV-1 drug resistance mutation prediction using quasispecies reconstruction supported analysis"

**Details of quasispecies reconstructed from clinical samples and their phylogenetic analysis**

| **Dataset** | **Protease** | **Reverse Transcriptase** | **Integrase** |
| --- | --- | --- | --- |
| D1 | 5 | 4 | 3 |
| D2 | 6 | 6 | 9 |
| D3 | 4 | 3 | 3 |
| D4 | 2 | 3 | 4 |
| D5 | 3 | 7 | 7 |
| D6 | 4 | 5 | 2 |
| D7 | 4 | 6 | 4 |
| D8 | 4 | 2 | 3 |
| D9 | 8 | 6 | 4 |
| D10 | 5 | 4 | 5 |
| D13 | 2 | 2 | 1 |
| D14 | 5 | 3 | 4 |
| D15 | 4 | 5 | 5 |
| D16 | 4 | 4 | 8 |
| D17 | 2 | 3 | 1 |
| D19 | 2 | 3 | 1 |
| D20 | 3 | 2 | 2 |
| D21 | 5 | 4 | 2 |
| D22 | 2 | 2 | 1 |
| D23 | 3 | 3 | 3 |

**Number of quasispecies in the clinical datasets**

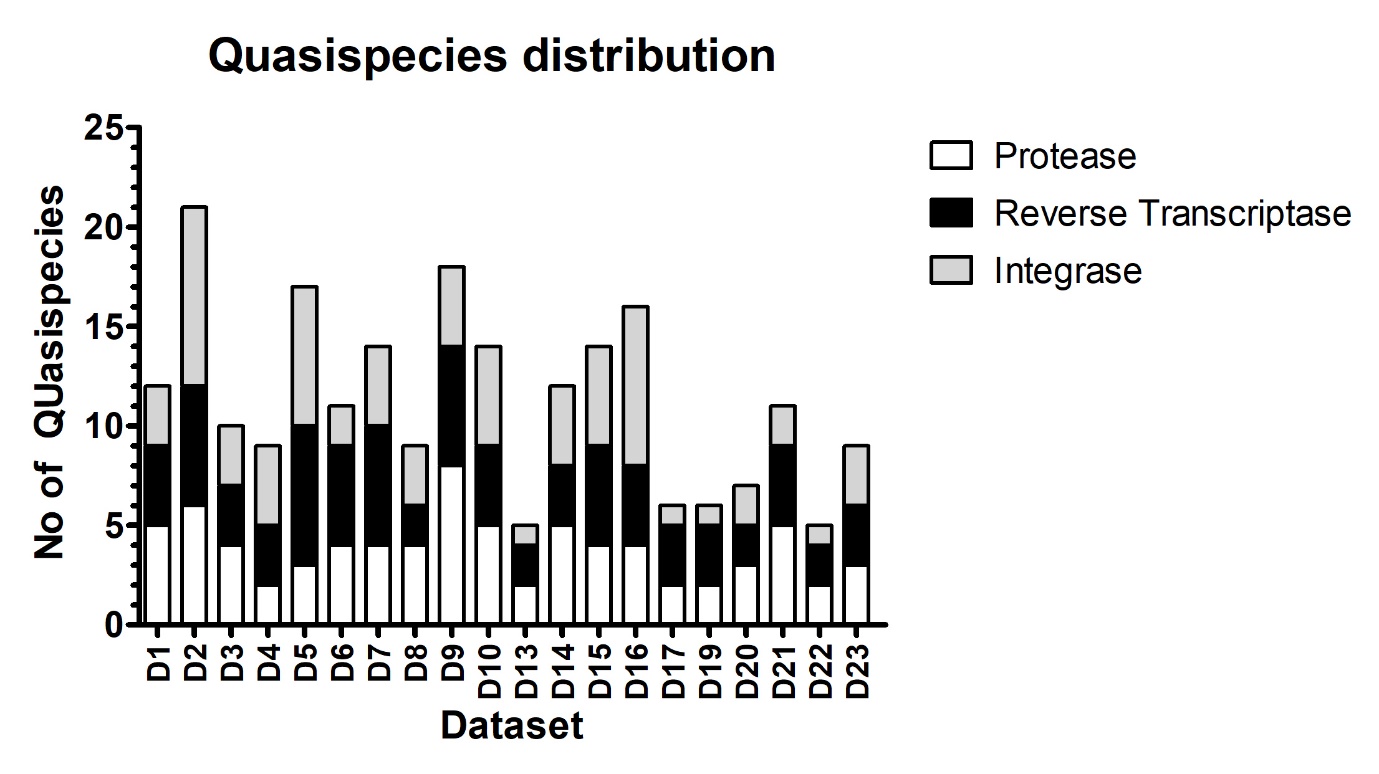

**Phylogenetic analysis of PR and RT genes**

Individuals with shared HIV transmission history have been color coded as D16: blue, D17: green. D21: red; D13: sky blue and D14: pink

**Protease gene**

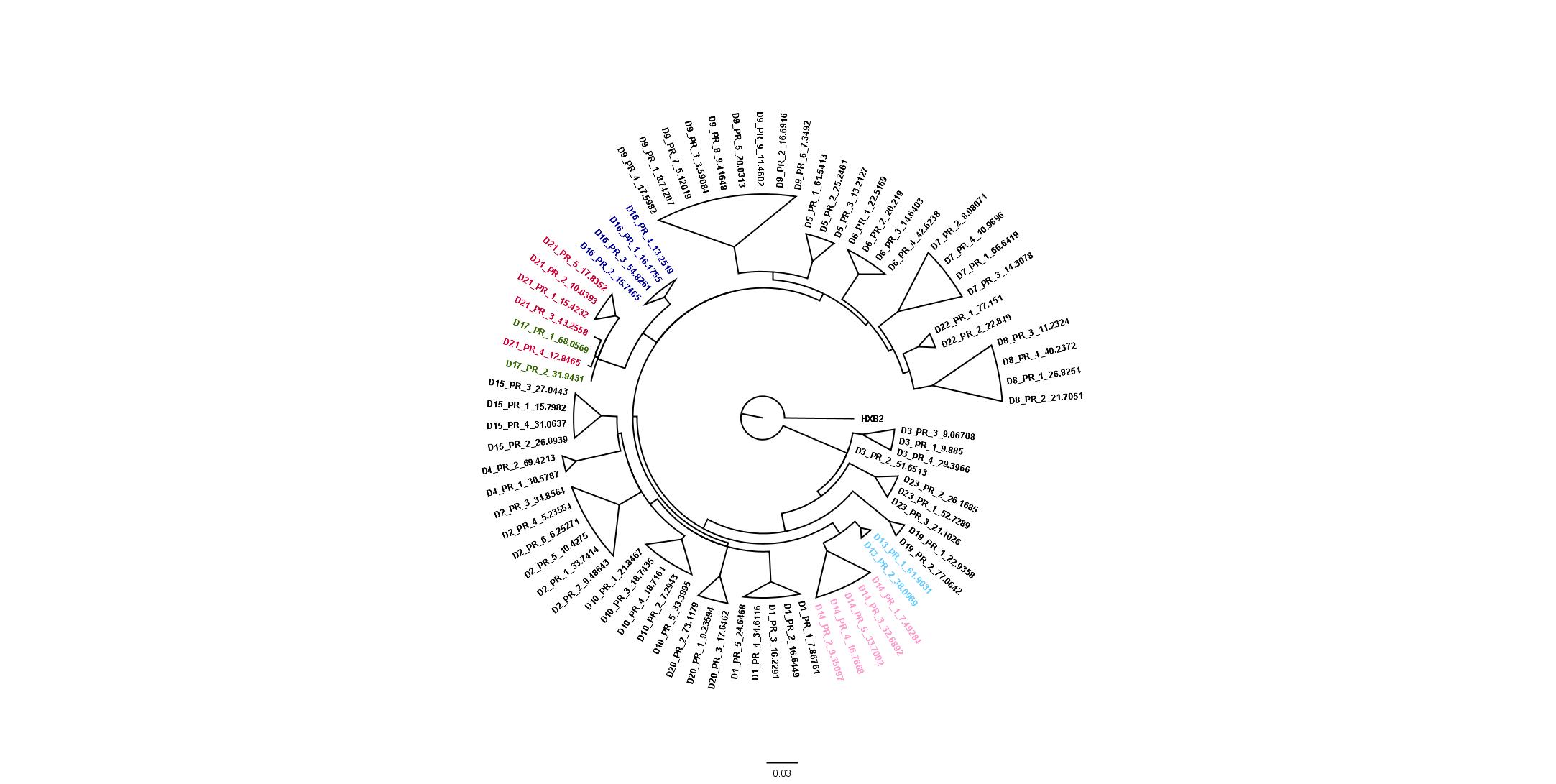

**Reverse transcriptase**

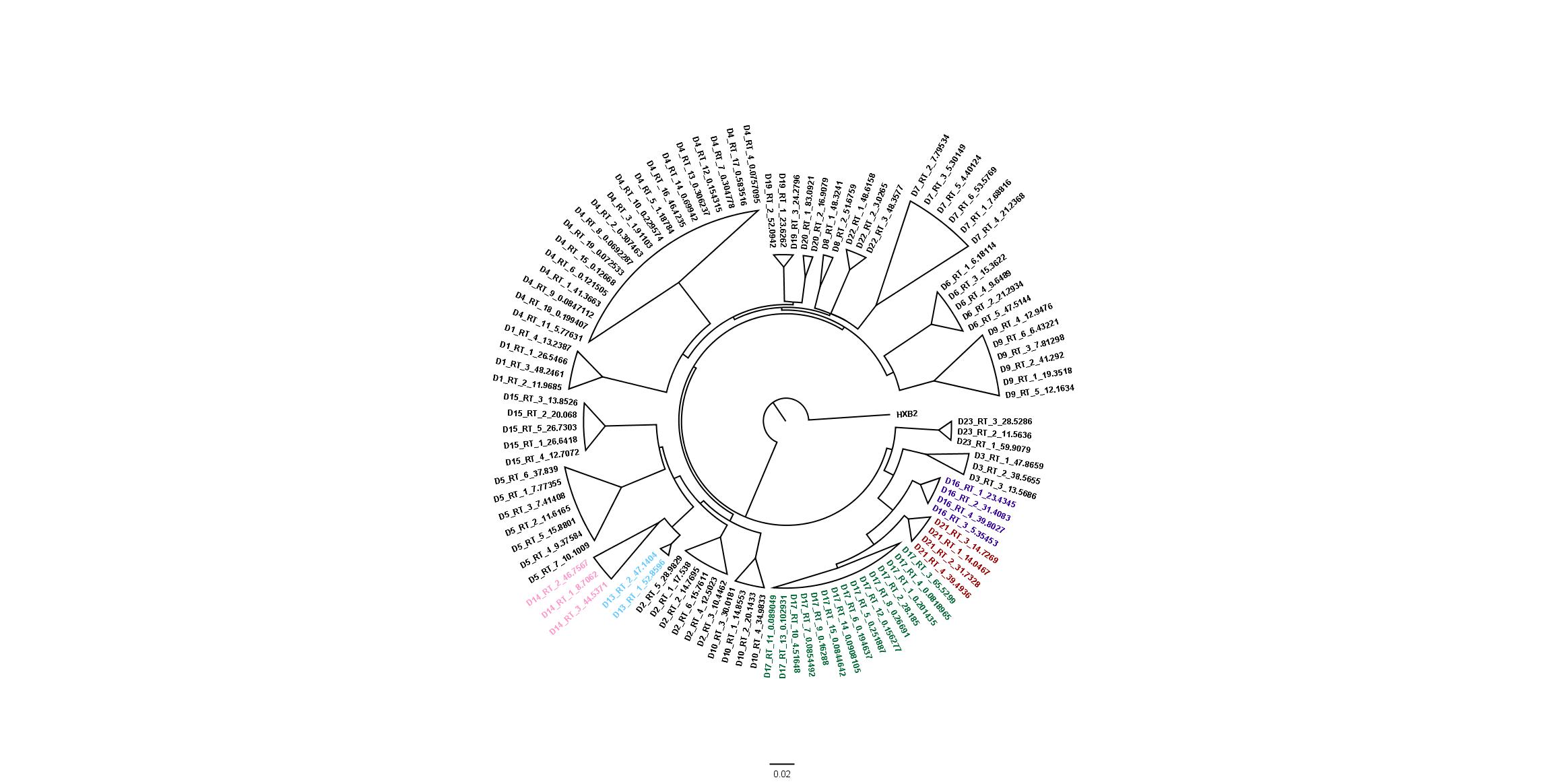
